# Investigation of the heterogeneity of cancer cells using single cell Ca^2+^ profiling

**DOI:** 10.1101/2024.01.14.575608

**Authors:** Amélie E. Bura, Camille Caussette, Maxime Guéguinou, Dorine Bellanger, Alison Robert, Mathilde Cancel, Margot Lacouette-Rata, Gaëlle Fromont, Christophe Vandier, Karine Mahéo, Thierry Brouard, David Crottès

## Abstract

Calcium (Ca^2+^) is an essential second messenger that controls numerous cellular functions. Characteristics of intracellular Ca^2+^ oscillations define Ca^2+^ signatures representatives of the phenotype of a cell. Oncogenic functions such as migration, proliferation or resistance to chemotherapy have been associated with aberrant Ca^2+^ fluxes. However, the identification of Ca^2+^ signatures representatives of the oncogenic properties of cancer cells remains to be addressed. To characterize oncogenic Ca^2+^ signatures, we proposed an unbiased scalable method that combines single cell calcium imaging with unsupervised and supervised machine learning tools. Here, we applied this method to investigate the heterogeneity of Ca ^2+^ signatures in 16 cancer cell lines and the remodeling of oncogenic Ca^2+^ signatures induced by acquired docetaxel resistance and in the course of the interaction of cancer cells with fibroblasts. Finally, our method demonstrates the potential of Ca^2+^ profiling for discriminating cancer cells and predict their phenotypic characteristics at single cell level, and provides a framework for researchers to investigate the remodeling of the Ca^2+^ signature during cancer development.

## Introduction

Calcium (Ca^2+^) is an essential and ubiquitous second messenger that controls numerous cellular functions (proliferation, apoptosis, migration, metabolism, etc.) and has been implicated in many pathologies including cancer. Ca^2+^ signaling relies on the finely tuned oscillations of the cytosolic Ca^2+^ concentration induced by components of the Ca^2+^ signaling toolkit (ion channels, pumps and ion exchangers, …) in response to input signals (Brodskiy & Zartman, 2018; Berridge *et al*, 2003). The spatio-temporal regulation of these cytosolic Ca^2+^ oscillations, the so-called “Ca^2+^ signature”, is essential to define the appropriate cellular response (Berridge *et al*, 2000). Different phenotypes, cell types or tissues will exhibit different characteristics of cytosolic Ca^2+^ oscillations (duration, frequency, amplitude, …) (Smedler & Uhlén, 2014; Boulware & Marchant, 2008; Evans & Blackwell, 2015; Gryshchenko *et al*, 2018). This heterogeneity of Ca^2+^ responses allows for the complexity of patterns associated with Ca^2+^ signaling, contributing to the diversification of cellular responses (Yokota *et al*, 2015; Scheuss *et al*, 2006).

Dysregulation of proteins that regulate Ca^2+^ signaling has been associated with various pathologies, including cancer (Marchi *et al*, 2020; Terrié *et al*, 2019; Guéguinou *et al*, 2014; Romito *et al*, 2022). Ca^2+^ fluxes have been linked to cancer hallmarks such as migration, proliferation or chemotherapy resistance (Marchi *et al*, 2020; Monteith *et al*, 2017; Chen *et al*, 2019; Figiel *et al*, 2019). Interestingly, the aberrant expression of ion channels is heterogeneous and depends on the type of cancer or the aggressiveness of the tumor (Chen *et al*, 2019; Bruce & James, 2020; Perrouin-Verbe *et al*, 2019). However, if the Ca^2+^ signature of cancer cells reflects this heterogeneous expression of Ca^2+^ signaling toolkit remains to be investigated.

Defining a Ca^2+^ signature is a complex task. Different individual descriptors (amplitude, duration, number of oscillations, latency, …) have been proposed to quantify and classify Ca^2+^ oscillations (Bootman *et al*, 2013). With recent advances in computational analysis, machine-learning based approaches have been proposed to analyze Ca^2+^ oscillations profiles. Unsupervised machine learning (kMeans) methods cluster neurons according to their Ca^2+^ oscillation profiles (Swain *et al*, 2018; Wang *et al*, 2024) and advanced deep learning algorithms identify patterns of Ca^2+^ oscillations in the cortex and suprachiasmatic nucleus associated with locomotor function and circadian rythms, respectively, in rodents (Wang *et al*, 2024; Ajioka *et al*, 2024). In recent years, machine- and deep-learning algorithms have been applied to MRI / CT images, sequencing data or histological H&E slides of various cancers and have demonstrated good accuracy for tumor cell detection and prediction of the aggressiveness (Zhong *et al*, 2024; Tran *et al*, 2021; Chuang *et al*, 2021; Cuocolo *et al*, 2019; Xu *et al*, 2024). With these methods, it becomes possible to handle massive amounts of data and to identify features associated to tumor development. However, these approaches collect data at discrete timepoints, and are therefore limited to assess cancer cell activity. A dynamic assessment of Ca^2+^ signaling in cancer cells could overcome this limitation and provide a novel approach to functionally characterize and identify therapeutic strategies target those cells.

To characterize and investigate the heterogeneity of oncogenic Ca^2+^ signatures, we proposed an unbiased scalable method that combines single cell calcium imaging with unsupervised and supervised machine learning tools. From an initial dataset of 27,439 agonist-induced Ca^2+^ responses elicited in a panel of 16 prostate and colorectal cancer cell lines, we discriminate 26 clusters of Ca^2+^ responses using unbiased unsupervised clustering inspired by single-cell transcriptomic workflows (Satija *et al*, 2015). From these clusters, we generate Ca^2+^ signatures for each cancer model. In parallel, we propose supervised neural network models predicting characteristics of a single cancer cell based on its profile of Ca ^2+^ responses. These approaches allow to investigate the remodeling of Ca^2+^ signatures in various pathological contexts. Here, we applied those methods to characterized a remodeling of Ca ^2+^ signatures associated with acquired docetaxel resistance or in the course of the interaction of cancer cells with fibroblasts. At single cell level, our supervised neural network succeeded to identify docetaxel-resistant cancer cells and to distinguish fibroblasts from cancer cells.

Taken together, our study proposes a new method to functionally investigate the cancer cell heterogeneity at the single cell level. Our study demonstrates the potential of Ca^2+^ profiling for discriminating cancer cells and predict their phenotypic characteristics, and provides a framework for researchers to investigate the remodeling of the Ca^2+^ signature during cancer development.

## Results

### Calcium (Ca^2+^) signaling pathway is aberrantly regulated in multiple cancers

Aberrant expression of proteins associated with Ca^2+^ signaling pathways has been reported in several cancers (Marchi *et al*, 2020; Monteith *et al*, 2017). Here, we performed a survey of 16 different cancers using transcriptomic data from The Cancer Genome Atlas (TCGA) consortium (https://portal.gdc.cancer.gov/). Compared to non-cancerous tissues, the gene set associated to the “KEGG Ca^2+^ signaling pathway” is significantly misregulated in primary tumors derived from 10 cancer types **(Fig. S1a)**. The differential expression of genes associated with the Ca^2+^ signaling toolkit in tumors is highly heterogeneous **(Fig. S1b)**. In colon and prostate cancers, both characterized by a downregulation of the Ca^2+^ signaling pathway, we observed distinct expression patterns of genes associated with this pathway (Abeshouse *et al*, 2015; Muzny *et al*, 2012) (**Fig. S1c**). Taken together, this suggests that despite the common misregulation of Ca^2+^ signaling pathways across cancer types, cancer cells may exhibit heterogeneous expression profiles of the Ca^2+^ signaling toolkit, which may be translated into heterogeneous functional Ca^2+^ signals representative of their origin or characteristics.

### Single cell Ca^2+^ imaging of cancer cell lines

To test this hypothesis, we proposed to define Ca^2+^ signatures of prostate (PCa) and colorectal (CRC) cancer cell lines using an unbiased approach combining single cell Ca^2+^ imaging, unsupervised and supervised machine learning algorithms (**Fig 1**.). Following the Depmap project, we selected cancer cell lines representative of the transcriptomic profiles of prostate and colon adenocarcinomas (**Fig. S1d**) (Warren *et al*, 2021). Each cell line was described by the cancer of origins (CRC or PCa), the site of isolation (non-cancerous tissues, primary tumor, or metastatic tumors) and the EMT score (epithelial or mesenchymal) computed using the Kolmogorov-smirnov method (Mandal *et al*, 2021). Cancer cells are non-excitable cells also it is not expected to observe spontaneous Ca^2+^ transients in these models. To have an overview of the variety of Ca^2+^ responses elicited by cancer cells, we propose to evaluate agonist-induced Ca^2+^ responses. We assayed for a panel of agonists whose receptors interact with proteins associated with the Ca^2+^ signaling toolkit. This allows the Ca^2+^ signaling toolkit to be stimulated in a variety of ways (**Fig. S1e**). Selected agonists included Adenosine Tri-phosphate (ATP), Lysophosphatidic Acid (LPA), Acetylcholine (Ach.), Platelet-Aggregating Factor (PAF), Histamine (Hist.), Epidermal Growth Factor (EGF), and Prostaglandin E_2_ (PGE_2_). Thapsigargin (Tg) and the vehicle (Veh.) were used as positive and negative controls, respectively, for the Ca ^2+^ response. Aside from Thapsigargin, all of the agonists have been shown in various models to influence Ca^2+^ signaling and impact key tumor phenotypes, such as proliferation and migration [ATP: (Shukla *et al*, 2024), Ach: (Bele *et al*, 2024), LPA: (Takai *et al*, 2024), PFA: (Tsoupras *et al*, 2009), EGF: (Crottès *et al*, 2019), PGE2: (Santiso *et al*, 2024), Histamine: (Azimi *et al*, 2024)]. Some of these agonists are also implicated in inflammatory cancers and/or play central roles in tumor development. All 16 cell lines were stimulated with each of 8 agonists. A total of 27,439 single cell Ca^2+^ responses were acquired and processed (**Fig S2**).

**Figure 1.**
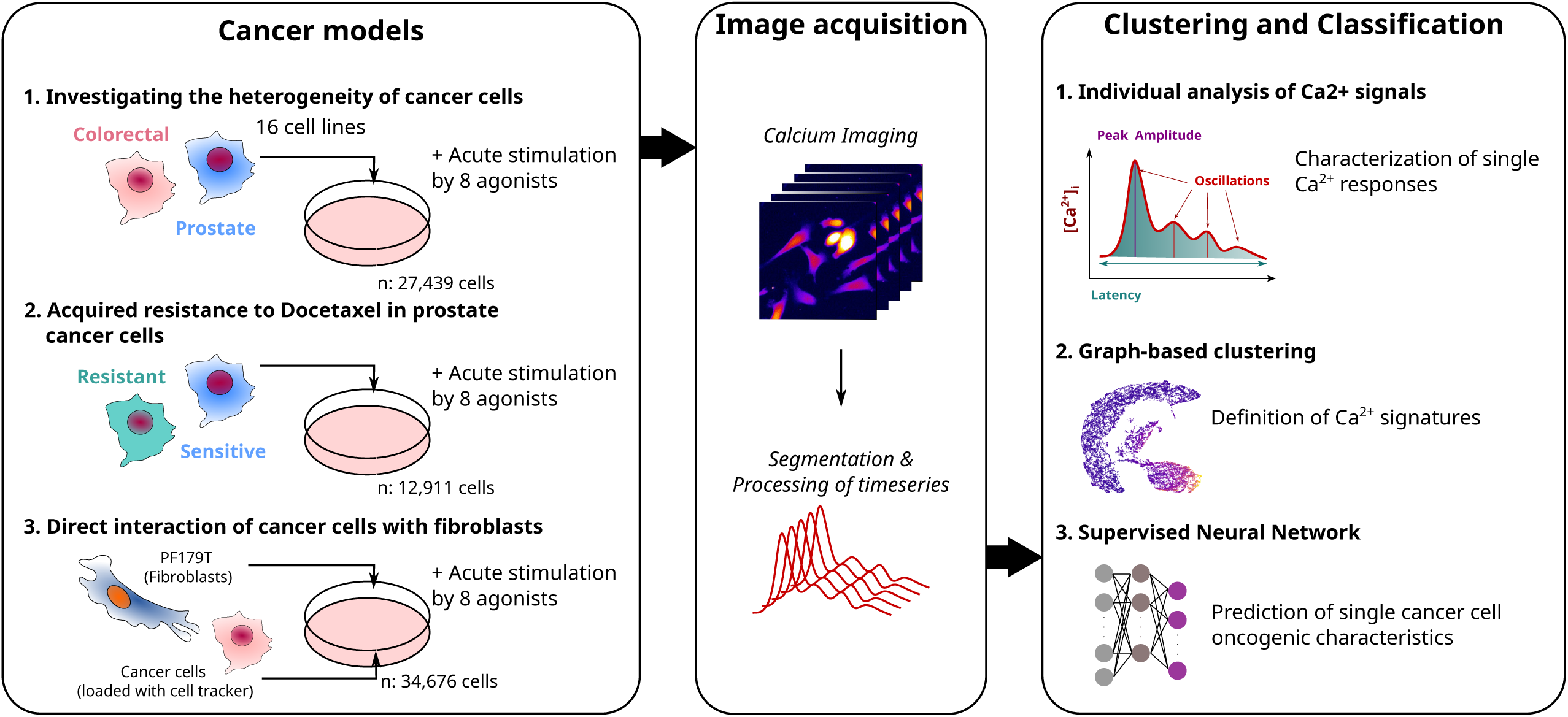
Heterogenous dys-regulation of Ca^2+^ signaling pathways in cancers. Schematic representation depicting the workflow of single cell Ca^2+^ profiling. Cells are loaded with fluorescent Ca^2+^ dye Cal-520TM. Agonist-induced Ca^2+^ responses of different cancer models are captured using time-lapse fluorescence microscopy. From each recording, individual cell fluorescence intensity are extracted by applying background correction and cell segmentation. Analysis of Ca^2+^ profiling is defined by computing 1) individual descriptors of Ca^2+^ responses are computed (peak amplitude, latency of responses, altered basal level, number of oscillations, percentage of responsive cells) from single cell Ca^2+^ response, 2) defining Ca^2+^ signatures of each cancer model by using an unsupervised clustering approach, 3) Assaying the prediction of oncogenic characteristics at single cell level using supervised classifier.

Over the years, a variety of parameters have been proposed to analyze and quantify Ca ^2+^ signals (Bootman *et al*, 2013). Considering the diversity and complexity of the acquired signals, we proposed here to limit our analysis to the description of the peak amplitude, the latency, the altered basal level, the number of oscillations and the proportions of responding cells (Bootman *et al*, 2013). The values obtained showed a large variability but a relatively uniform distribution (**Fig S3**). When comparing cancer cells according the cancer of origins (CRC vs PCa), the site of isolation (Primary, Metastatic or non-cancerous tissues) or the EMT Score (Epithelial vs Mesenchymal), we observed similar distributions of values of each parameter **(Fig S4 and S5)**. This large heterogeneity of agonist-induced Ca^2+^ responses makes it difficult to define a Ca^2+^ signature based on a few parameters. To circumvent these problems, we proposed to analyze the whole signal using unsupervised machine learning tools and propose an unbiased clustering of the profiles of Ca^2+^ responses.

### Unsupervised clustering of heterogeneous agonist-induced Ca^2+^ responses identified Ca^2+^ signature of cancer cell lines

In analogy to the clustering workflow used in single-cell RNA-Seq (Satija *et al*, 2015; Patterson-Cross *et al*, 2021), we developed an approach that sequentially reduced the number of dimensions by principal component analysis (PCA), built a shared-nearest-neighbor network on the first 20 components, and clustered the nodes of the network using a Louvain algorithm (Blondel *et al*, 2008) (**Fig 2a and Fig S6**). This approach identified 5 groups of Ca^2+^ responses subdivided into 26 clusters (**Fig 2b-c** and **Fig S7**). Each cluster and group of clusters gathers single cell Ca^2+^ responses with particular characteristics. Thus, clusters of Ca^2+^ responses associated with groups A and B are from unresponsive cells (**Fig 2d-e**). Clusters associated with group E gather single cell Ca^2+^ response with the highest peak amplitude and number of oscillations. Clusters of the group C and D are associated with the highest value of the altered basal level and of the latency respectively (**Fig 2d-e**).

**Figure 2.**
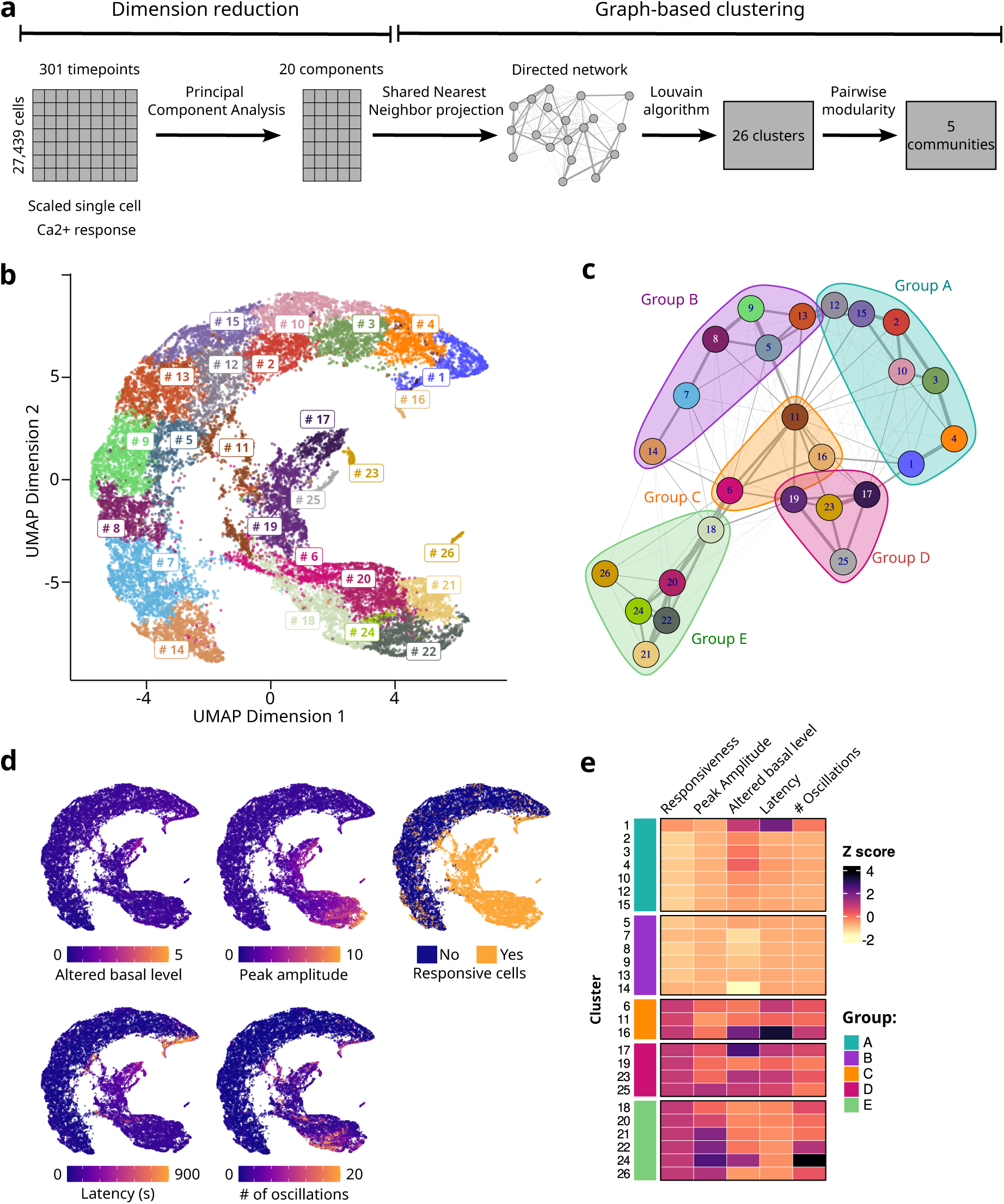
Unsupervised clustering of agonist-induced single cell Ca^2+^ responses. **a.** Schematic workflow representing the unsupervised clustering of agonist-induced single cell Ca^2+^ response. **b.** Uniform Manifold Approximation and Projection (UMAP) representation of 27,439 agonist-induced single cell Ca^2+^ responses. Each dot represents a single cell and colored according 26 clusters defined by Louvain community detection algorithm applied on shared nearest neighbor graphs (SNN). **c.** Force-directed layout graph representing cluster modularity highlighting five communities of clusters. **d.** UMAP representation of single cell Ca^2+^ responses color-coded according values of Altered basal level, peak amplitude, responsiveness, latency, and numbers of oscillations. **e.** Heatmap representing the relative value of responsiveness, peak amplitude, altered basal level, latency, and number of oscillations of single cell Ca ^2+^ responses averaged by clusters.

We defined Ca^2+^ signatures by observing the distribution of single cell Ca^2+^ responses elicited by each agonist in each cluster for each cancer models (**Fig 3a**). We examined the similarity of these Ca^2+^ signatures across cell lines by principal component analysis and computing the Wasserstein distance (also called Earth Mover’s distance or EMD, a pairwise measure of the dissimilarity of two-dimensional objects) (**Fig 3b-c**).

**Figure 3.**
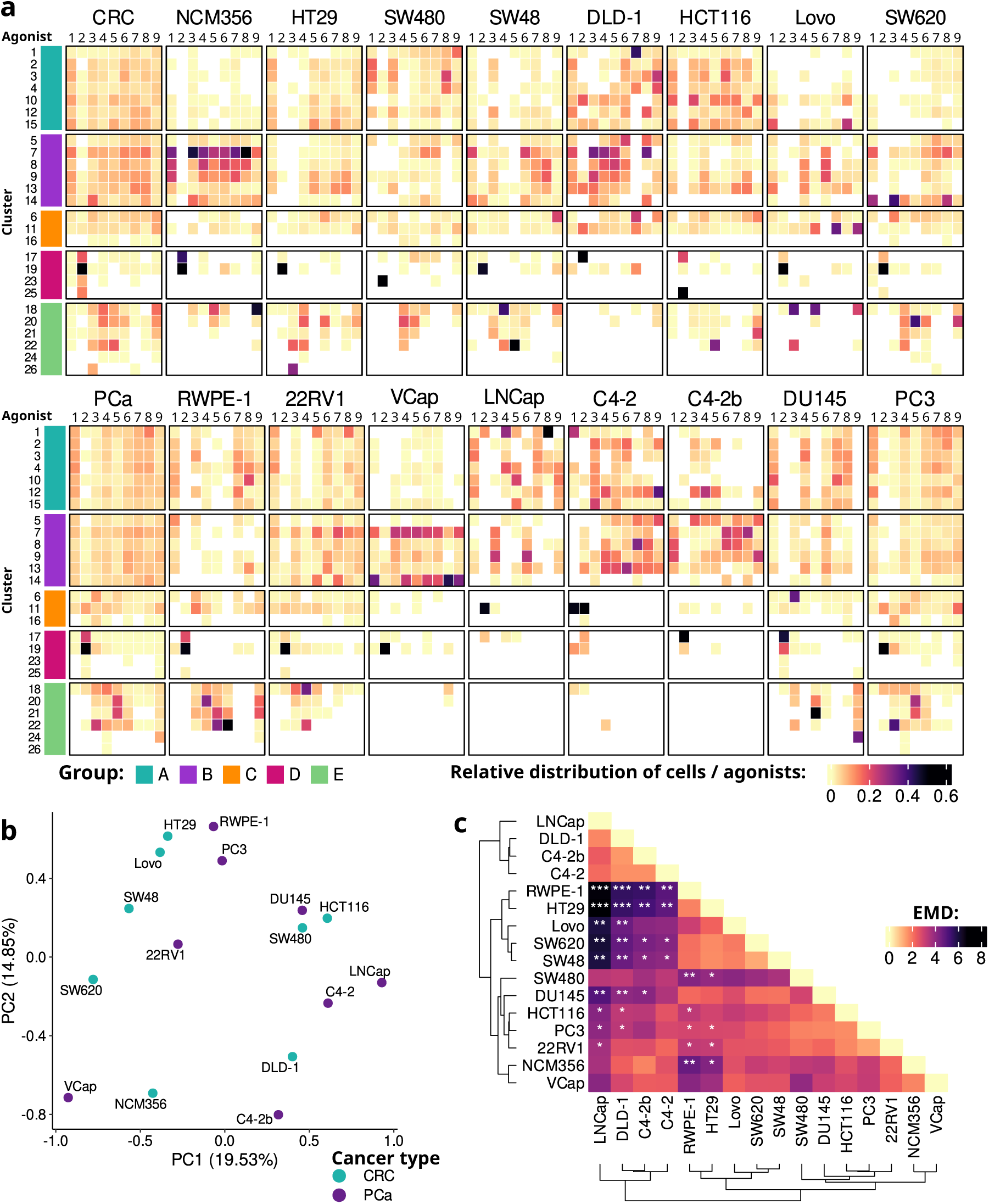
Definition of Ca^2+^ fingerprints of individual cancer cell lines. **a.** “Ca^2+^ fingerprint” of each cell line represented by the relative proportion of agonist-induced Ca^2+^ responses associated to each cluster. **b.** Principal components (PC) analysis of cell lines according the Ca^2+^ fingerprint. c. Pairwise measure of the Wasserstein distance (Earth mover’s distance or EMD) representing the dissimilarity between Ca^2+^ fingerprints. (Significant p values are calculated by permutation test: * p < 0.05, ** p < 0.01 and *** p < 0.001). Agonists are: 1- vehicle (Veh.), 2- Thapsigargin (Tg), 3- Adenosine Tri-phosphate (ATP), 4- Acetylcholine (Ach.), 5- Lysophosphatidic Acid (LPA), 6- Platelet-Aggregating Factor (PAF), 7- Epidermal Growth Factor (EGF), 8- Prostaglandin E2 (PGE2), 9- Histamine (Hist.).

Comparison of the average Ca^2+^ signature for CRC and PCa cell lines revealed a relatively similar pattern (**Fig 3a**). This was confirmed by principal component analysis of the Ca^2+^ signatures of each cell line, which showed a mixed spatial distribution of cell lines according to their cancer of origin (**Fig 3b**). Dissimilarity matrix analysis revealed that cell lines clustered into three groups with distinct Ca^2+^ signatures (**Fig 3c**). None of these groups are associated with a single cancer type suggesting a redundancy in the profile of Ca^2+^ responses across cancers. However, it is interesting to note that cell lines with similar genetics such as C4-2, C4-2b and their parent cell line, LNCap are grouped together and show similar Ca^2+^ signatures. We can also note that the SW480 and SW620 cancer cell lines have different Ca^2+^ signatures, even though these cell lines were isolated from primary and metastatic colon tumors, respectively, from the same individual. This may suggest that primary tumors evolving as metastasis will be subject to a remodeling of their Ca^2+^ signatures. However, we did not observe a significant dissimilarity when comparing Ca^2+^ signatures of cancer cell lines isolated from primary or metastatic sites (**Fig S8**). No significant difference was observed either between Ca^2+^ signatures of epithelial and mesenchymal cancer cells or between cancer cells isolated from primary / metastatic tumors and non-cancerous tissues (**Fig S8**).

Taken together, graph-based clustering of agonist-induced Ca^2+^ responses defines Ca^2+^ signatures of cancer cell lines. While we did not identify significant difference between Ca ^2+^ signatures of cancer cells according their cancer type, their site of isolation or their EMT score, this approach allows to classify cancer cell lines according their Ca^2+^ signatures and to highlight some similarity or dissimilarity between cancer models. This may indicate a certain degree of redundancy in the profile of Ca^2+^ responses induced in different cancers.

### The agonist-induced Ca^2+^ response of a single cell is predictive of its phenotype

Ca^2+^ fingerprints show that each cancer cell line has a specific Ca^2+^ signature. We then wondered whether the Ca^2+^ response profile elicited by a single cell might reflect its characteristics and be associated with its lineage or cancer of origin. To verify this hypothesis, we developed a three-layer artificial neural network (ANN) that was trained to predict the cell lineage associated with a single cell agonist-induced Ca^2+^ response (**Fig 4a** and **Fig S9a,d**). We obtained different results depending on the class (cell lineage) predicted. The model had the best performance for cells derived from the PC3 cell line (F1-score: 0.722±0.017) and the worst for those derived from the LNCap cell line (F1-score: 0.211±0.068) (**Fig 4b and Fig S9b**). We then trained the ANN to predict the cancer type (2 classes: CRC or PCa) (**Fig 4c**). Similar values were obtained for the F1-score, with 0.814±0.003 and 0.805±0.01 for the prediction of CRC and PCa cells respectively (**Fig 4d**). The AUROC values for both classes were close to 0.9, indicating good class discrimination by our model (**Fig S9c**). Recall of the model for predicting of each class (CRC or PCa) as a function of the agonist-induced Ca^2+^ responses showed that ATP and Tg-induced Ca^2+^ responses had the best predictive performance for both classes. We can also note that Ach., LPA and PAF-induced Ca^2+^ responses have significantly different predictive performance for each cancer type (**Fig S9e**). Calculation of Shapley values (SHAP) shows that the first and last timepoints, and the timepoints immediately after agonist addition (timepoint 20 corresponding to 1 minute of the recording) have the highest weight in predictive performance (**Fig S9f**). Similarly, this supervised approach generate a high F1 score when trained to identify the EMT score or the site of isolation (primary / metastatic tumors) of a single cell according its agonist-induced Ca^2+^ response (**Fig 4d-e**).

**Figure 4.**
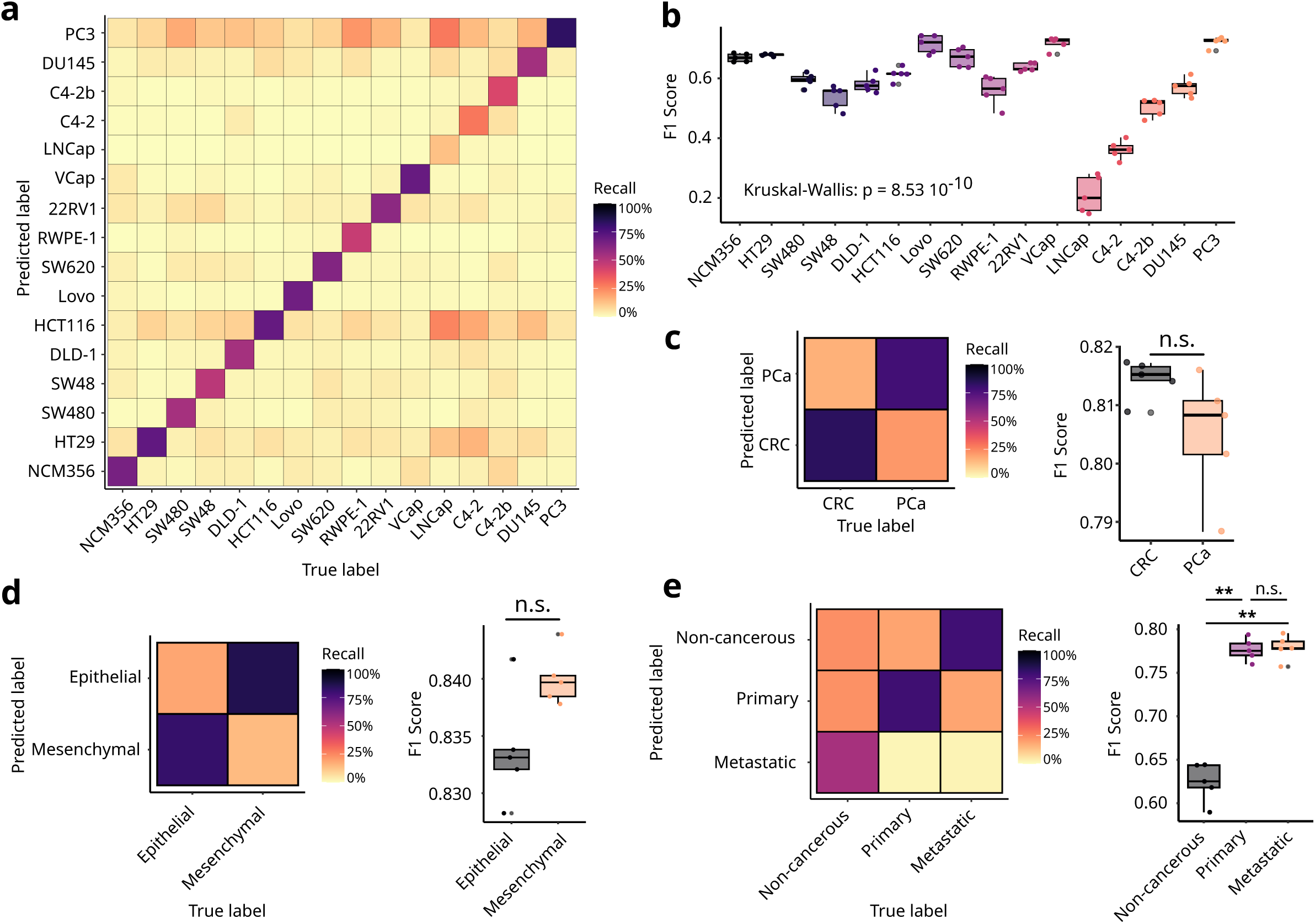
Predicting single cell identity using artificial neural network trained with Ca ^2+^ responses. Artificial neural network (ANN) model was trained for multi-class classification of single cell Ca^2+^ responses according their lineage or their cancer type. Input are the values of each timepoint of the measure of Ca^2+^ response and the label of agonist used. Each model are trained with 80% of the dataset for training and 20% for validation. **a. A**verage confusion matrix of 5 train / validation settings representing the recall of prediction of lineage. **b.** Boxplot representing the F1 score of the prediction of each lineage. **c.** Average confusion matrix of 5 train / validation settings of the ANN representing the recall of prediction of the cancer of origin of each cell. Boxplot representing the F1 score of the prediction of each cancer type. **d.** Prediction of the site of isolation of cancer cell lines (Non-cancerous tissues, Primary or Metastatic tumors) by ANN. Average confusion matrix (Left) and F1 score (right) of 5 train / validation settings representing the recall of prediction of site of isolation. **e.** Average confusion matrix (Left) and F1 score (right) of 5 train / validation settings representing the recall of prediction of the EMT score of each cancer cell. (Significant p values are calculated by Kruskal-wallis (b.) or Mann-Whitney test to compared “validation” and “test” set (c-e): * p < 0.05, ** p < 0.01 and *** p < 0.001).

Thus, we demonstrated that supervised machine learning models can successfully predict the cancer type or the lineage associated with a single cell using its agonist-induced Ca ^2+^ response as input.

### Acquired chemoresistance induces a remodeling of Ca^2+^ signature

Under therapeutic pressure, tumors often develop molecular mechanisms of resistance to treatment. Proteins associated with Ca^2+^ signaling have been implicated in the development of these features (Romito *et al*, 2022). Here, we investigate whether acquired chemoresistance induces a remodeling of the Ca^2+^ signature of cancer cells and whether we can identify resistant cancer cells at the single cell level.

We developed docetaxel-resistant 22RV1 and PC3 cell lines. For PC3 (IC50 for docetaxel: 0.66±0.18 nM), we used two different protocols to generate 3- and 30-fold more resistant cell lines (designated PC3R^Low^ and PC3R^High^, respectively, with IC50 for Docetaxel of 2.33±0.317 nM and 19.2±8.42 nM). Docetaxel-resistant 22RV1 (referred to as 22RV1R) is 10-fold more resistant than its parental cell line (IC50 of 22RV1 and 22RV1R for docetaxel are respectively of: 1.99±0.431 nM and 20.7±4.22 nM, respectively) (**Fig 5a**). We observed that the distribution of single cell Ca^2+^ responses associated with docetaxel-resistant cell lines into clusters previously identified by our unbiased approach was slightly modified, generating distinct Ca^2+^ signatures (**Fig 5b, c** and **Fig S10a-c**). Pairwise dissimilarity analysis revealed that the Ca^2+^ signature of PC3R^High^ is significantly modified compared to its sensitive counterpart (**Fig 5d**). PC3R^Low^ has a similar Ca^2+^ fingerprint to the docetaxel-sensitive PC3, suggesting that moderate (3-fold) acquired resistance does not strongly alter the Ca^2+^ signature. Despite a high EMD value, the Ca^2+^ fingerprint of 22RV1R is not significantly different from that of 22RV1 (p value = 0.068).

**Figure 5.**
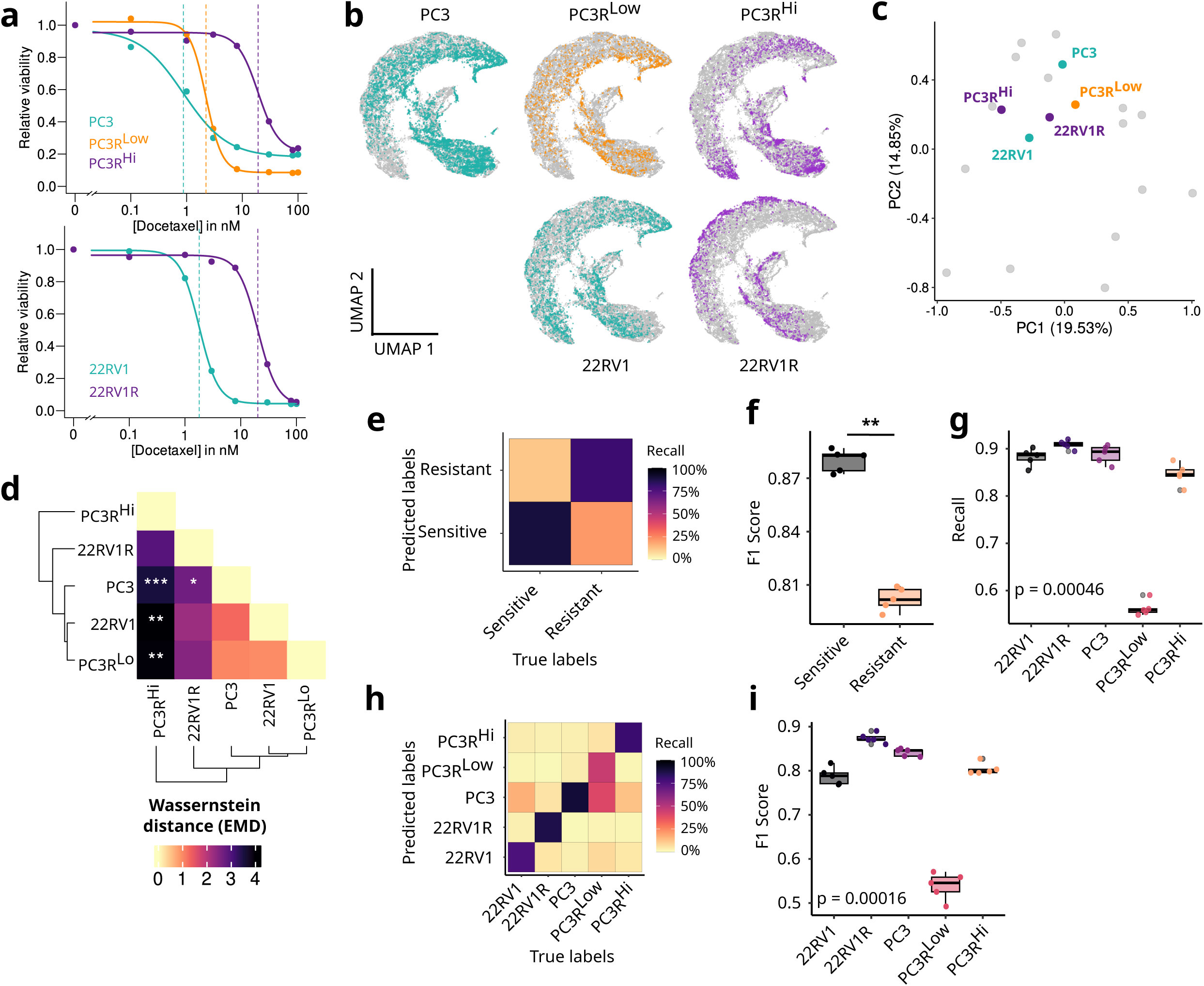
Investigating Docetaxel induced resistance in prostate cancer cell lines using unsupervised Ca^2+^ profiling. **a.** Cell viability of Docetaxel resistant PC3RHigh, 22RV1R and moderately resistant PC3RLow cell lines and respective parent PC3 and 22RV1 cell lines. **b.** UMAP projection of single cell agonist-induced Ca^2+^ responses of PC3, PC3RLow, PC3RHigh, 22RV1 and 22RV1R. Grey points represent single cell Ca^2+^ responses from 16 different cell lines depicted in Figure 2a. **c.** PC projection of Ca^2+^ footprints of PC3, PC3RLow, PC3RHigh, 22RV1 and 22RV1R. Grey points represent Ca^2+^ footprints of 16 different cell lines depicted in Figure 2d. **d.** Pairwise measure of EMD between global Ca^2+^ footprints of PC3, PC3RLow, PC3RHigh, 22RV1 and 22RV1R. (Significant p values are calculated by permutation test: * p < 0.05, ** p < 0.01 and *** p < 0.001). **e.** Average confusion matrix of 5 train / validation settings of the ANN representing the recall of prediction to the resistance to Docetaxel. **f-g.** Boxplot showing the F1 score **(f)** or the recall of each cell line **(g)** for predicting resistance to Docetaxel. h. Average confusion matrix showing the recall of ANN trained to predict sensitive and Docetaxel resistant sub-cell lines. **i.** Boxplot showing the F1 score for predicting sensitive and Docetaxel-resistant subcellular lines. (Significant p values are calculated by Kruskal-wallis (g-i.) or Mann-Whitney test (f.): * p < 0.05, ** p < 0.01 and *** p < 0.001).

We then investigated whether single cell Ca^2+^ responses could predict the resistance status of these cell lines. We trained the ANN with 12,911 single cell Ca^2+^ responses elicited in 22RV1, 22RV1R, PC3, PC3R^Low^ and PC3R^High^ (**Fig S10a-b**). The model successfully predicted the resistance status associated with each single cell (**Fig 5e**). We noticed a drop in performance when predicting resistant cells (**Fig 5f**), which we attributed to the poor prediction performance when confronted with single cell Ca^2+^ responses of PC3R^Low^ (**Fig 5g**). This may be explained by the similarity of the Ca^2+^ signatures of PC3 and PC3R^Low^. We trained our ANN to predict the cell line associated with each single cell Ca^2+^ response. As expected, our model showed poor performance in predicting PC3R^Low^, misattributing most of the PC3R^Low^ samples to PC3 (**Fig 5h** and **5i**).

Thus, we have identified a change in the Ca^2+^ signature of cancer cells that acquire resistance to chemotherapy. Our model has some limitations in identifying moderately resistant cell lines compared to sensitive cells. However, the Ca^2+^ response profile of a single cell seems to be a good indicator to classify sensitive and resistant cancer cells.

### Direct co-culture with fibroblasts induces a remodeling of the Ca^2+^ signature of cancer cells

The interaction of cancer cells with the microenvironment is fundamental for tumorigenesis. Surrounding non-cancer cells such as fibroblasts, adipocytes, or immune cells can influence the phenotype of cancer cells. It has been observed that interactions between cancer cells and cancer-associated fibroblasts can modify the Ca^2+^ signaling pathway of cancer cells (Sadras *et al*, 2021).

Here, we tested whether our supervised model is able to discriminate between non-cancerous and cancerous cells in a mixed environment. We co-cultured cancer cells with fibroblasts (cancer-associated fibroblast cell line PF179T). Cancer cells were preloaded with a red fluorescent cell tracker (Red CMPTX). After 72h, we acquired agonist-induced Ca^2+^ responses (**Fig 6a** and **Fig S11a-c**). We evaluated the performance of the ANN to discriminate cancer cells from fibroblasts using single cell Ca^2+^ responses from both cell types cultured alone or in co-culture. The model was trained with 34676 single Ca^2+^ responses. We observed high F1 scores for both cell types (fibroblasts: 0.830±0.004, cancer cells: 0.864±0.01) (**Fig 6b-c**). We observed that the recall of cancer cell prediction is significantly lower when the model is confronted with single cell Ca^2+^ responses originated from cancer cell lines co-cultured with fibroblasts (**Fig 6d**). Regarding the prediction of fibroblasts, PF179 co-cultured with HCT116 and PC3 shows a lower recall of prediction than PF179T culture alone (**Fig 6e**). These results suggest that while our model successfully distinguishes cancer cells from fibroblasts, the co-culture conditions affect the Ca^2+^ signature of cancer cells, resulting in a decrease in performance for their prediction. We confirmed this by comparing the Ca^2+^ signatures of cancer cells cultured alone or in the presence of fibroblasts (**Fig 6f**). Pairwise dissimilarity analysis revealed that PC3 Ca^2+^ signatures were significantly altered by the presence of PF179T (**Fig 6g**). HCT116 and HT29 Ca^2+^ signatures are not significantly modified by their co-culture with PF179T. Similarly, no significant differences were observed between PF179T cultured alone or in the presence of cancer cells (**Fig 6g**).

**Figure 6.**
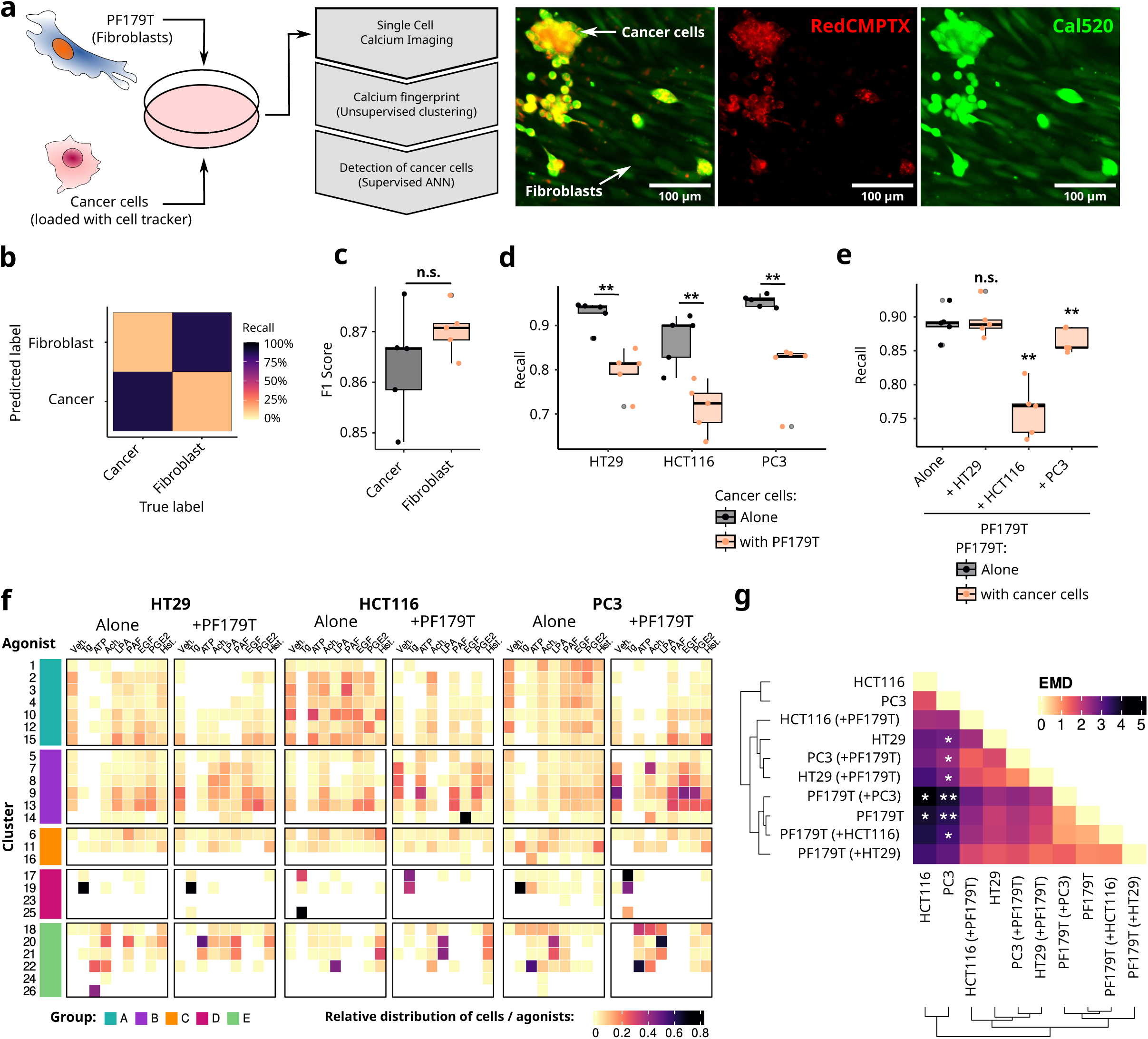
Interaction of cancer cells and fibroblasts induces a remodeling of Ca^2+^ signature. **a.** Model of co-culture between cancer cells / fibroblasts. Cancer cells are loaded with RedCMPTX are cultured with PF179T for 72h prior single cell Ca^2+^ imaging. On the left, representative fluorescent images of cancer cell / fibroblasts cultured. **b.** ANN trained with 34,676 single cell Ca^2+^ responses of PF179T, HT29, HCT116 and PC3 cultured alone or with cancer cells / fibroblasts. Confusion matrix representing the recall of prediction of cancer cells / fibroblasts. **c.** Boxplot representing the F1 score of the prediction of Cancer cells / Fibroblasts. **d.** Boxplot representing the performance (recall) for the classification of cancer cells of single cell Ca^2+^ responses of each cancer cell lines cultured alone or in presence of PF179T. **e.** Boxplot representing the performance (recall) for the identification of fibroblasts using single cell Ca ^2+^ responses of PF179T cultured alone or in presence of cancer cells. **f.** “Ca^2+^ fingerprint” of cancer cell line cultured alone or in presence of PF179T represented by the relative proportion of agonist-induced Ca^2+^ responses associated to each cluster. **g.** Pairwise measure of the Wasserstein distance (Earth mover’s distance or EMD) representing the dissimilarity between Ca^2+^ fingerprints. The name in parenthesis indicate the cell line used for cocultured with the cell line compared. (Significant p values are calculated by Man-Whitney test **(c-e)** permutation test **(g)**: * p < 0.05, ** p < 0.01 and *** p < 0.001).

Taken together, these data suggest that single cell Ca^2+^ response can be an indicator to distinguish cancer cells from surrounding non-cancerous cells. The Ca^2+^ signatures are modulated by the interaction of cancer cells with their microenvironment, providing a functional window to study the crosstalk between intra-tumoral compartments.

## Discussion

In this study, we present two machine-learning based methods using Ca^2+^ signaling to characterize and investigate cancer cell heterogeneity. First, graph-based unsupervised clustering of agonist-induced Ca^2+^ responses defines the Ca^2+^ signature associated with multiple cancer cell lines and allow functional comparison of different models. Second, we demonstrate that our neural network models can identify individual cancer cells and predict their characteristics (origin, aggressiveness, lineage, sensitivity to chemotherapy) from the profile of the Ca^2+^ response suggesting that this intracellular signal carries information representative of the identity and phenotype of the cell.

Using these methods, we investigate the remodeling of Ca^2+^ signaling in docetaxel-resistant cancer cell lines and during the course of cancer cell / fibroblast interaction. We find that acquired chemoresistance induces a remodeling of the Ca^2+^ signature of prostate cancer cell lines. This is consistent with previous reports describing the overexpression of various ion channels involved in Ca^2+^ signaling in chemoresistant cancer cells (Romito *et al*, 2022). Interestingly, we observed that cancer cells can be distinguished from fibroblasts by their Ca ^2+^ response profile and that the interaction of both cell types induces a remodeling of their respective Ca^2+^ signatures. Our results are thus in line with previous reports describing the influence of the stroma on the expression of proteins involved in the Ca^2+^ signaling pathway in cancer cells (Sadras *et al*, 2021).

Overall, our study provides the proof of concept that single cell Ca^2+^ profiling can distinguish cancer cells and identify key characteristics. It provides a framework to investigate cancer cell heterogeneity at the functional level, thus offering a new tool to study the consequences of cancer cell evolution on both the tumor and the micro-environment compartment.

Single cell calcium profiling has the advantage of functionally assessing the cellular identity, thus complementing other genomic, transcriptomic, and proteomic approaches. In contrast to these methods, single cell calcium profiling is relatively inexpensive and can be performed on any fluorescence microscope. Additionally, it does not require genetic engineering and can be applied to virtually any cell line (primary or immortalized).

In conjuction with single cell calcium imaging, we propose two machine learning modules to facilitate the integration and analysis of Ca^2+^ profiles. First, a graph-based unsupervised clustering approach inspired by the single-cell transcriptomic workflow (Satija *et al*, 2015) allows the definition of Ca^2+^ signatures. The advantages of this module are that 1) it reduces the introduction of analytical bias by processing the entire signal rather than extracting a few descriptors, 2) it facilitates the scalability by using normalized and scaled signals as input, and 3) it provides a two-dimensional Ca^2+^ fingerprint (or signature) that can be qualitatively compared across conditions independently of the number of cells acquired. This overcomes similar approaches that rely on either the manual annotation or the definition of an arbitrary number of clusters of Ca^2+^ responses to be identified (James *et al*, 2011; Swain *et al*, 2018; Mohammed *et al*, 2017). In our study of 27,439 single cell agonist-induced Ca^2+^ responses, we identified 26 clusters of agonist-induced Ca^2+^ responses organized into 5 distinct communities (Fig 2b-c). This clustering highlights the diversity of patterns of Ca^2+^ responses elicited by cancer cells. It should be noted that this number of clusters is not absolute and likely depends on the type of agonist selected, its concentration and the cell line studied. Further studies with additional agonists / models should be carried on in order to verify this clustering of agonist-induced Ca^2+^ responses and/or to identify additional clusters of Ca^2+^ responses.

Second, we proposed a module that integrates artificial neural network models trained to predict either the origin, the cell type, the lineage or the sensitivity to docetaxel of a single cell from its profile of Ca^2+^ response profile. Each of these models provides high overall predictive performance. However, we note a relatively low predictive performance for some class labels (LNCap, C4-2, C4-2b cell lines or PC3R^Low^ cell line) which can be explained either by a general absence of Ca^2+^ responses elicited in these models or by a Ca^2+^ response profile relatively similar to that of the parent cell line. To our knowledge, this is the first time that a supervised machine learning approach has been used to define single cell identity using Ca^2+^ response profiles. The predictive performance of these relatively simple models supports the idea that Ca^2+^ signaling can define a cellular identity and be representative of its origin and its characteristics. It also demonstrates the potential of our approach to identify specific biomarkers related to cancer cell response to treatment, thereby strengthening its utility for precision oncology.

A potential limitation of our study is that the immortalized cell lines used may not recapitulate the full spectrum of tumor characteristics observed in the clinic, although they may retain similar genomic and transcriptomic profiles (Warren *et al*, 2021; Tang *et al*, 2022; Berg *et al*, 2017). In addition, we have here examined a panel of single-dose agonists whose receptors interact with the Ca^2+^ signaling toolkit in a variety of ways and known to be associated with oncogenic functions (Fig S1c). However this panel is limited and it is possible that additional agonists could induce unknown Ca^2+^ response profiles not captured in this study.

Single cell Ca^2+^ profiling is a valuable tool to characterize and investigate the functional activity of cancer cells. Future studies investigating Ca^2+^ signatures in tumors *in vivo* (or *ex vivo* using organotypic culture or patient-derived organoid models) will be interesting to 1) compare *in vivo* Ca^2+^ signatures with those observed *in vitro* and 2) provide a functional overview of the organization of different cellular compartments in and around the tumor. Here, we observe in a direct co-culture model of cancer cells and fibroblasts, we observe that each cell type exhibits distinct Ca^2+^ signatures opening the possibility to map individual cell types on tumor tissues at functional level using Ca^2+^ profiling. Consistent with this, a difference in the amplitude of Ca^2+^ fluxes has been reported when comparing organotypic cultures obtained from prostate tumors and non-cancerous tissues (Figiel *et al*, 2019). Recent studies have used convolutional neural networks to decode Ca^2+^ signals from the mouse cortex and suprachiasmatic nucleus and correlate these signals with walking function and circadian rhythms (Wang *et al*, 2024; Ajioka *et al*, 2024). Using this method, they identified a distinct brain region involved in the Ca^2+^ signature associated with these functions. Taken together, Ca^2+^ profiling could represent a valuable tool to spatially and functionally annotate the multicellular organization of tumoral tissues.

Overall, the single cell Ca^2+^ profiling workflow presented in this study is a high throughput, scalable platform that considerably expand our ability to investigate Ca^2+^ signaling, its heterogeneity and improves our understanding of the coding cellular response by Ca ^2+^ signaling. In cancer research, this study provides a proof of concept that cancer cells can be defined by their Ca^2+^ signature and conversely that Ca^2+^ signaling can serve as a predictor of cancer cell characteristics. Further studies are needed to evaluate Ca^2+^ signatures *in vivo*, to investigate their remodeling during tumor development and to determine whether targeting these functional signatures can influence cancer cell phenotype.

## Materials and Methods

### Cell culture

The colorectal cancer (CRC) cell lines (HT29, DLD1, HCT116, Lovo, SW48, SW480, and SW620) were cultured in McCoy media supplemented with GlutaMax (Gibco, Thermo Fisher, Illkirch, France), 10% fetal bovine serum (FBS; Eurobio, Les Ulis, France) and 1% penicillin and streptomycin (Eurobio). NCM356 cells (Incell Corporation, LLC, San Antonio, TX, USA) were cultured in high-glucose Dulbecco’s Modifed Eagle Medium (DMEM) (Sigma-Aldrich, Missouri, USA) supplemented with 10% FBS and 1% penicillin and streptomycin (Eurobio). Prostate cancer cell lines 22RV1, DU-145, C4-2 and C4-2b were cultured in RPMI-1640 media + GlutaMax (Gibco, Thermo Fisher, Illkirch, France) supplemented with 10% fetal bovine serum (FBS; Eurobio, Les Ulis, France) and 1% penicillin and streptomycin (Eurobio). PC3 were cultured in RPMI-1640 media + GlutaMax (Gibco, Thermo Fisher, Illkirch, France) supplemented with 5% fetal bovine serum (FBS; Eurobio, Les Ulis, France) and 1% penicillin and streptomycin (Eurobio). VCap cell line was cultured in DMEM supplemented with 2 mM Glutamine (Eurobio), 10% fetal bovine serum (FBS; Eurobio, Les Ulis, France) and 1% penicillin and streptomycin (Eurobio). Fibroblast cell line PF179T was cultured in α-MEM supplemented with 10% FBS and 1% penicillin and streptomycin. All cells were cultured in an incubator at 37°C with 5% CO_2_ and do not exceed 15 passages.

### Generation of Docetaxel-resistance cell lines

Docetaxel-resistant 22RV1R and PC3R^High^ cell lines were generated by stepwise increased concentrations of docetaxel (Sigma-Aldrich, Missouri, USA) over a period of 6 months (starting from 0.5 nM to 4 nM). Docetaxel moderately resistant PC3R^Low^ cell lines were generated by repeated intermittent treatment of docetaxel (2 nM) (72h of treatment followed by 72h without treatment) over a period of 8 weeks.

### Single cell calcium Imaging

Single cell calcium imaging was performed as described previously (Crottès *et al*, 2019). Each day of experiments include one positive control (such as Tg or LPA) per cell lines tested. Briefly, 48 h before the experiment, 5×10^4^ cells were plated on 35-mm dishes (Fluorodish, WPI, USA). For cell density experiments, 5×10^4^, 15× 10^4^, or 30×10^4^ cells were plated on 35-mm dishes. For co-culture experiments, 5×10^4^ of PC3, HCT116, HT29 or NCM356 pre-loaded with red cell tracker dye (RedCMPTX, Invitrogen, USA) at 5 µM for 45 min were mixed with 4× 10^5^ PF179T cells per 35 mm dish. Cells were then cultured for 72h in alpha-MEM supplemented with 10% FBS and 1% penicillin and streptomycin. On the day of the experiment, cells were loaded with Cal520-AM (4.5 µM, AAT Bioquest, Pleasanton, USA) at 37°C for 30 min in physiological saline solution (PSS) containing: 150 mM NaCl, 4 mM KCl, 1 mM MgCl_2_, 2 mM CaCl_2_, 10 mM Hepes, and 10 mM D-glucose. Cells were then washed twice in PSS and incubate in PSS solution for 3 min before acquisition. For stimulation of cells, an equal volume of a 2× concentrated compound of interest (prepared in PSS) was added to the dish after the first minute of acquisition. Fluorescence was acquired using either a Nikon TiEclipse inverted scope equipped with a DS-Qi2 camera (Nikon, Tokyo, Japan), and a SOLA SE-5-LCR-SA Lamp (Photometrics, Germany), or a Nikon Ti2 inverted scope equipped with a Fusion BT camera (Hamamatsu) and a pE-800 (CoolLed, United Kingdom). A combination of dichroic cube, excitation and emission filters at 488 and 515 nm (SemRock, USA) filter fluorescence emitted by Cal520-AM, Images were acquired every 3s for 15 min.

### Calcium imaging analysis

All videos were processed using a custom macro on Fiji / ImageJ software (NIH, Bethesda, USA). Images were subjected to rolling background substraction. Automated cell segmentation was performed using Cellpose plugin (Pachitariu & Stringer, 2022) on maximal projection of timeseries and manually checked. Fluorescence intensity of individual cells was then measure and normalizing to the fluorescent intensity measured in the first image (F/F_0_).

### Processing of calcium profiling

All profiles of calcium measurement were processed using R 4.4.1 and Rstudio 2023.03.0 Build 386. Multiple cell batches for the majority of cell type-agonist pair have been used. No major difference in the distribution of patterns of Ca^2+^ responses were observed for each batch. For those that were tested only once, the absence of multiple batches represents a limitation that as to be taken in consideration. For each single cell, maximal amplitude, time to reach the maximal amplitude and AUC were calculated after the first minute of acquisition. Cells are considered as positively responsive for the tested compounds if the fluorescence is superior to 110% of the initial fluorescence (F_0_). Oscillations were determined using *find_peaks* functions from *GCalcium* R package (n.points = 3). Only oscillations with a fluorescence above 110 % of the initial fluorescence (F_0_) are counted.

### Dimensionality reduction (UMAP)

For visualization in two dimensional spaces, single cell Ca^2+^ responses are normalized and subjected to Uniform Manifold Approximation and Projection (UMAP) using *umap* R package. n_neighbors and min_dist parameters were modulated to assay for the best visualization. Unless otherwise is stated, min_dist = 0,1 and n_neighbors = 15 were used for UMAP representation.

### Graph-based Unsupervised clustering

Each single cell calcium response are normalized and scaled. Principal component analysis (PCA) was applied on scaled dataset and first 20 components are extracted for clustering. Unsupervised clustering was performed using *bluster* and *igraph* R packages. Briefly, a shared-Nearest Neighbor (SNN) graph network is built by applying a *J*accard weighting scheme on reduced scaled dataset. Louvain algorithm for community detection identified clusters on the SNN graph. Pairwise modularity associated to Louvain clustering highlights group of clusters sharing similar values. Optimal values (k=15 and resolution = 1.2) were defined empirically by evaluating the segregation of calcium responses in each cluster. For further analysis of additional single cell Ca^2+^ responses, a k-Nearest-Neighbors (kNN) classifier is trained using the initial clustering provided for 27,439 single cell Ca ^2+^ responses to predict the cluster associated to each additional single cell Ca^2+^ responses.

### Supervised Machine learning and Neural Networks

Artificial Neural network (ANN) was built using *tensorflow, keras, scikit-learn* and *numpy* Python packages. Architecture of ANN models is composed of three layers of neurons and use Softmax activation function and Adam optimiser. The first layer is the *input* layer containing normalized and scaled fluorescence intensity recorded at each timepoints and binary values for each agonist, 2) *C1 “hidden”* layer of half-sized the input layer followed by a dropout layer (coefficient: 0.4) to reduce overfitting and 3) *output* layer composed of different possibles labels. Unless otherwise stated, default training:validation ratio of 0.8:0.2 was applied on the dataset. Mean squared error was used as a loss function. Learning rate and batch size were set at 9e−6 and 32 respectively. ANN were trained for 500 epochs and for each models, 5 different runs were performed. Performance were evaluated by calculating F1 score, AUROC recall and accuracy. For evaluating performance of the model for predicting the cancer type of an “unknown” cell line, train / validation were performed on data obtained from 15 cell lines. Testing were performed using calcium responses measured on the 16^th^ cell line.

### Explainability of neural network

Shap values were determined by using shap python packages. Briefly, 1000 randomly selected samples were used as background dataset to use for integrating out features. Then, shap values were estimated for 500 samples for each prediction class and representing the contribution of each features to the model output. Shap values are then represented as relative absolute values for quantitative features (timepoints) and absolute values for qualitative features (agonists). Importance of each features in the predicted labels of the validation test were alternatively evaluated by permutation and F1 score were re-calculated.

### Transcriptomic analysis

Transcriptomic data from 16 different cancer types were generated by The Cancer Genome Atlas Research Network (https://portal.gdc.cancer.gov/). Read counts were downloaded from GDC using the *tcgaworkflow* package and used for differential expression analysis (DEA) using the package *edgeR*. DEA compared non-cancerous and primary tumors for each cancer types. The ranked list of genes differentially expressed according their log2(fold change) was submitted to Gene Set Enrichment Analysis (GSEA) using the fgsea package and gene set collections from MSigDB (https://www.gsea-msigdb.org/gsea/msigdb-c2.5.go.bp.v7.5.1.symbols.gmt). Alignement of transcriptomic profiles of cancer cell lines with primary and metastatic tumors were obtained from DepMap project (https://depmap.org/portal/) (Warren *et al*. 2021). EMT score of each cell line were computed using the two-sample Kolmogorov Smirnov test as described previously (Mandal *et al*, 2021).

### Viability assay

Cell viability of Docetaxel-sensitive and resistant 22RV1 and PC3 cell lines were evaluated using standard MTT assay after 7 days of treatment. Briefly, cells were treated with MTT for 1h at 37°C and then formazan crystals were dissolved using DMSO over 10 min. Absorbance at 570 nm was determined with a BioTek (Vermont, United States) Spectrophotometer.

### Statistical analysis and illustration

Figures are generated using either *ggplot2*, *alluvial*, *ComplexHeatmap* R packages and assembled using Inkscape software. Statistical test (Kruskall-Wallis and post-Dunn test, Wilcoxon test and ANOVA) were performed using *ggpubr* and *rstatix* R packages. Dissimilarity index for matrices Earth Mover’s distance (also called as Wassernstein distance) was computed using *emddist* package.

### Datasets and code availability

The microscopy calcium imaging data reported in this paper will be shared by the lead contact upon request. Datasets used in this paper are deposited on Figshare (doi: 10.6084/m9.figshare.28202924). R and Python scripts used for unsupervised clustering and neural networks are added as supplemental files.

## Acknowledgments

We acknowledge all lab members of Inserm U1069 and LIFAT for their help. We thank Dr. Chin Fen Teo and Dr. Sami Tuomivaara for their helpful comments. This work was supported by the Le Studium, Loire Valley Institute for Advanced Studies, Orléans & Tours, France (DC), Région Centre-Val de Loire APR-IR LCAPRO (CC, AB and KM) and APR-IA CAMITHERAPAL (MG) grants, Inserm PCSI MULTINIR (CV), Cancéropôle Grand-Ouest IMACALSEQCAP (DC) and LNCRESIST grants (KM and DC), ARFMAD association (MG) and the following french department committees of Ligue Contre le Cancer “Grand-Ouest” (KM and DC): 56 (Morbihan), 41 (Loir et Cher), 18 (Cher). The funders had no role in study design, data collection and analysis, decision to publish, or preparation of the manuscript.

