## Supplementary material for "Investigation of the heterogeneity of cancer cells using single cell Ca^2+^ profiling": Suplemental data

**SUPPLEMENTAL MATERIAL FOR: Investigation of the heterogeneity of cancer cells using single cell $Ca^{2+}$ profiling.**

#### EXTENDED MATERIALS AND METHODS:

**Artificial neural network – scripts and output.**

Artificial neural networks used in this study were all designed following the same design organized around 4 steps:

1- Data preparation:

Dataset 3 including scaled single cell $Ca^{2+}$ response is used. Column "Agonist" is transformed from long to wide format, encoded as a binary description for each possible output (1 = presence, 0 = absence) and joined to timepoints of scaled fluorescence (columns "0" to "300) and named as "X" dataframe. Columns "ID_Cancer", and "Cell Lines" are transformed from a long to a wide format and encoded as a binary description for each possible output (1 = presence, 0 = absence) and iteratively used as prediction output (named "y").

2- Building of the model:

A 3-layer ANN model is built using *keras* and *tensorflow* python libraries. Briefly, the first layer "input" is composed of as many neurons as the number of values of input (X and Y). Second layer "C1" is composed of half the number of neurons than the first layer. A dropout layer (coefficient: 0.4) is added to prevent overfitting. Third layer "output" is composed of the same number of neurons as desired output. First and second layers applied the rectified linear unit activation (ReLU) on data and the third and final layer applied Softmax activation function

3- Training of the model:

X and Y dataframe are splitted into a training (X_train and Y_train) and testing set (X_test and Y_test) at a ratio of 0.8 / 0.2. X_train is scaled using *StandardScaler* function and the same sacling function is apply independently to X_test. ANN model is then trained using X_train and Y_train during 500 epochs and *Adam* and *mean squared error* as functions for respectively optimizer and loss. Trained model is saved in "model" folder and values of loss and accuracy for each epoch are save in "history" folder.

4- Testing the model:

Performance of the model is evaluated using X_test and Y_test. Matrix confusions are saved in "conf_mat" folder. F1 score, AUROC, TPR and FPR values are saved in "roc_auc_f1". Shap values are calculated using a subset of 1000 samples of X_train and saved in "shap" folder. For evaluating the importance of each feature in the prediction, values of each feature (column) of X_test are randomly permuted and results of matrix confusion, F1 score, AUROC are saved in "Predictions_values" folder.

**Supplementary Figures Legends:**

**Supplemental Figure 1:**

**a.** Normalized enrichment score of the « KEGG $Ca^{2+}$ signaling pathway » gene set in 16 different cancer types from the The Cancer Genome Atlas (TCGA) public repository. Dotted lines indicate a non-significant NES value. **b.** Heatmap showing the log fold change between tumor and non-cancerous samples of each gene associated with the $Ca^{2+}$ signaling pathway in 16 different cancers. **c.** Heatmap highlighting the heterogeneous expression of normalized log mRNA expression of genes associated with the $Ca^{2+}$ signaling pathway in TCGA-COAD (colon cancer) and TCGA-PRAD (prostate cancer). **d.** UMAP 2D projection of colon and prostate adenocarcinoma primary tumor (TCGA), metastatic tumor (Met500) samples and cell lines (DepMap) expression data according Celligner methods published by Warren et al. (2021). **e.** Force layout network projections showing the interactions of proteins of the $Ca^{2+}$ signaling toolkit with receptors and proteins interacting with Adenosine Tri-phosphate (ATP), Lysophosphatidic Acid (LPA), Acetyl-Choline (Ach.), Platelet-Aggregating Factor (PAF), Histamine (Hist.), Epidermal Growth Factor (EGF), Prostaglandin $E_2$ ($PGE_2$) and Thapsigargin (Tg). Interactions are obtained by String app on cytoscape software.

**Supplemental Figure 2.** Single cell $Ca^{2+}$ imaging of colon and prostate cell lines. Single cell $Ca^{2+}$ responses of a panel of CRC (in green) and PCa cell lines (in purple) has been recorded during 15 minutes. After one minute, differents ligands were added extracellularly. (ATP: Adenosine Tri-phosphate, Ach: AcetylCholine, EGF: Epidermal Growth Factor, LPA: Lysophosphatidic Acid, PAF: Platelet Aggregating Factor, PGE2: Prostaglandin E2, Tg: Thapsigaring, Veh.: Vehicle).

**Supplemental Figure 3.** Intra-lineage heterogeneity of single cell $Ca^{2+}$ responses. Density of the peak amplitude, the latency, the altered basal level, the number of oscillations and the percentage of responding cells in each cell lines stimulated by agonists.

**Supplemental Figure 4.** Heterogeneity of single cell $Ca^{2+}$ responses in function of the cancer type of the site of isolation of cancer cells. Density of the peak amplitude, the latency, the altered basal level, the number of oscillations and the percentage of responding cells for each cancer types (**a.**) or the site of isolation (**b.**).

**Supplemental Figure 5.** Heterogeneity of single cell $Ca^{2+}$ responses according the EMT score. Density of the peak amplitude, the latency, the altered basal level, the number of oscillations and the percentage of responding cells for epithelial and mesenchymal cancer cells stimulated by agonists.

**Supplemental Figure 6.** Optimization of parameters for unsupervised clustering of single cell $Ca^{2+}$ responses. A. Single cell $Ca^{2+}$ responses are clusterized by sequentially reduced the number of dimension by Principal Component Analysis (PCA), build a shared-nearest neighbor network (SNN) on top of first 20 components and clusterized nodes of the network using a Louvain algorithm. For the definition of the optimal parameters, variables values for the parameters numbers of neighbors (k) for SNN and the resolution of Louvain algorithm were assayed. Optimal configuration of unsupervised clustering were determined by parameters allowing a low average standard deviation of clusters and % of clusters with a standard deviation > 0.1. Additionally, high average silhouette width, purity and sum of squares per clusters were indicators of good performance for the unsupervised clustering. Here, we defined the optimal configuration by using k:37 and resolution: 1.2.

**Supplemental Figure 7.** Visual representation of clusters defined for each agonist-restricted single cell $Ca^{2+}$ responses. Average $Ca^{2+}$ responses per clusters are in red.

**Supplemental Figure 8:**

**a.** “$Ca^{2+}$ fingerprint” of epithelial and mesenchymal cancer cells determined by the EMT score.

**b.** “$Ca^{2+}$ fingerprint” of cells isolated from non-cancerous tissues, primary or metastatic tumors.

**Supplemental Figure 9:**

**a.** Architecture of artificial neural network (ANN) model for the prediction of each cell line and cancer types using as input the values of each timepoint of the measure of $Ca^{2+}$ response and the label of agonist used.

**b-c.** ROC curves representing the sensitivity and specificity of the prediction for each cell line (**b**) or cancer types (**c**). Inset: value of Area under the ROC curve (AUROC) for each.

**d.** Representation of the accuracy and loss for training and validation set during the training of the ANN.

**e.** Boxplot showing the recall of ANN for the true prediction of the cancer type of individual $Ca^{2+}$ response associated to a particular agonist.

**f.** Relative absolute SHAP value for each quantitative features (timepoints) of different ANN tested. Values are the mean of 5 independently trained ANN.

**Supplemental Figure 10:**

**a.** Agonist-induced single cell $Ca^{2+}$ response of sensitive (22RV1) and Docetaxel resistant (22RV1R) sub-cell lines

**b.** Agonist-induced single cell $Ca^{2+}$ response of sensitive (PC3) and Docetaxel moderately resistant ($PC3R^{Low}$) and resistant ($PC3R^{High}$) sub-cell lines.

**c.** "$Ca^{2+}$ fingerprint" of 22RV1 and PC3 sensitive ad docetaxel resistant cell lines.

**Supplemental Figure 11:**

**a.** UMAP projection of single cell agonist-induced $Ca^{2+}$ responses of PF179T cultured alone or in presence of HT29, HCT116 and PC3. Grey points represent single cell $Ca^{2+}$ responses from 16 different cell lines depicted in Figure 2a.

**b.** "$Ca^{2+}$ fingerprint" of PF179T cultured alone or in presence of HT29, HCT116 and PC3

**c.** UMAP projection of single cell agonist-induced $Ca^{2+}$ responses of HT29, HCT116, PC3 cultured alone or in presence of PF179T. Grey points represent single cell $Ca^{2+}$ responses from 16 different cell lines depicted in Figure 2a.

**d.** "$Ca^{2+}$ fingerprint" of HT29, HCT116, PC3 cultured alone or in presence of PF179T .

**Supplementary Dataset 1. Single cell $Ca^{2+}$ response.** Table containing all measure of the relative fluorescence ($F/F_0$) of the calcium dye Cal-$520^{AM}$ representative of the intracellular $Ca^{2+}$ concentration.

*"ID":* Unique ID for a single cell**,**

*"ID_Cancer" :* Designation of the cancer type (PCa: Prostate Cancer, CRC: Colorectal Cancer),

*<u>"Cell_Line":</u>* Code designation of the cell line used (NCM356, HT29, SW480, SW48, DLD-1, HCT-116, SW620, Lovo, RWPE-1, 22RV1, Vcap, LNCap, C4-2, C4-2b, DU145, PC3),

*<u>"Agonist":</u>* Code designation of the agonist used (Vehicle: "2Ca-2Ca", Thapsigargin (2 µM): "2Ca-Tg2uM", ATP (1 µM): "2Ca-ATP1uM", Acetylcholine (1 µM): "2Ca-Ach1uM", LPA (1 µM): "2Ca-LPA1uM", PAF (100 nM): "2Ca-PAF100nM", EGF (15 nM / 100 ng/mL): "2Ca-EGF100", PGE2 (50 µM): "2Ca-PGE5uM", Histamine (10 µM): "2Ca-Hist10uM"),

*<u>"Cell":</u>* Designation of cell ID per experiments,

"1": Relative fluorescent intensity measured at timepoint 0 (0 secondes),

"2": Relative fluorescent intensity measured at timepoint 1 (3 secondes),

…

"301": Relative fluorescent intensity measured at timepoint 300 (15 minutes)

**Supplementary Dataset 2. Computed parameters of single cell $Ca^{2+}$ response.** Table containing the value of parameters representative of $Ca^{2+}$ response. All values were computed after the timepoint 19 (1 minute of acquisition, time of injection of the agonist) to be representative of the agonist-induced $Ca^{2+}$ response.

- "Peak_Amplitude": Maximal value of fluorescence,
- "NumberOscillation": Number of oscillations defined by the function find_peaks of Gcalcium R package,
- "Responding": $Ca^{2+}$ response is considered positive if the amplitude of $Ca^{2+}$ response is beyond 110% of the initial fluorescence (F0),
- "Latency": Timepoint corresponding to the maximal amplitude of fluorescence,
- "Altered_basal": difference of fluorescence between last and first timepoints.

**Supplementary Dataset 3. Scaled single cell $Ca^{2+}$ response.** Values of each single cell $Ca^{2+}$ response from Dataset 1 were scaled using the formula: Scaled individual value = (individual value – minimal value of the dataset) / (maximal value of the dataset – minimal value of the dataset).

**Supplementary Dataset 4. Unsupervised clustering of scaled single cell $Ca^{2+}$ response.** Global or agonist-restricted unsupervised clustering was achieved using a Louvain algorithm on top of shared-nearest neighbor network representation. UMAP dimensionality reduction of $Ca^{2+}$ responses are computed to generate graphics presented in Figure 2B and Figure 3A and B.

- “Cluster”: designate the cluster ID obtain after unsupervised clustering of $Ca^{2+}$ responses.
- “Communities”: designate the Community ID obtain after cluster modularity.
- “UMAP_1”: Dimension 1 obtained after UMAP on $Ca^{2+}$ responses
- “UMAP_2”: Dimension 2 obtained after UMAP on $Ca^{2+}$ responses

**Supplementary Dataset 5. Scaled single cell $Ca^{2+}$ response of docetaxel resistance cell**.

*Docetaxel_resistance :* Indicate if a cell belong to a sensitive or docetaxel resistant cell line.

**Supplementary Dataset 6. Scaled single cell $Ca^{2+}$ response of the co-culture model.**

*“Cell_Line”:* Code designation of the co-culture model (PF179T+HT29, PF179T+HCT116, PF179T+PC3).

*Identified_cell_type :* Code designation for single cell indicating which cell lines (PF179T, HT29, HCT116, PC3). Values are defined based on fluorescence intensity of RedCMPTX.

### SUPPLEMENTARY FIGURE 1.

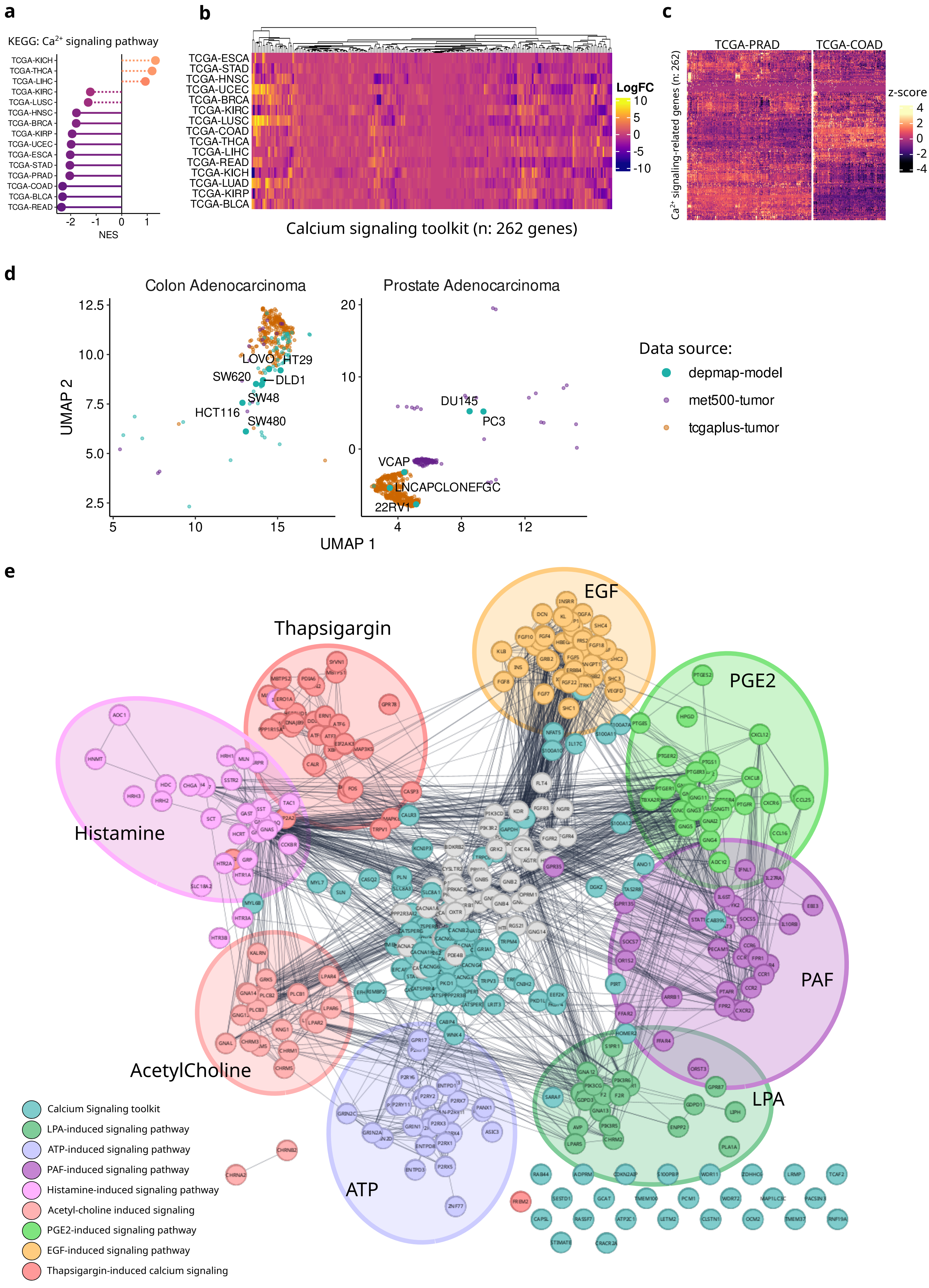

### SUPPLEMENTARY FIGURE 2.

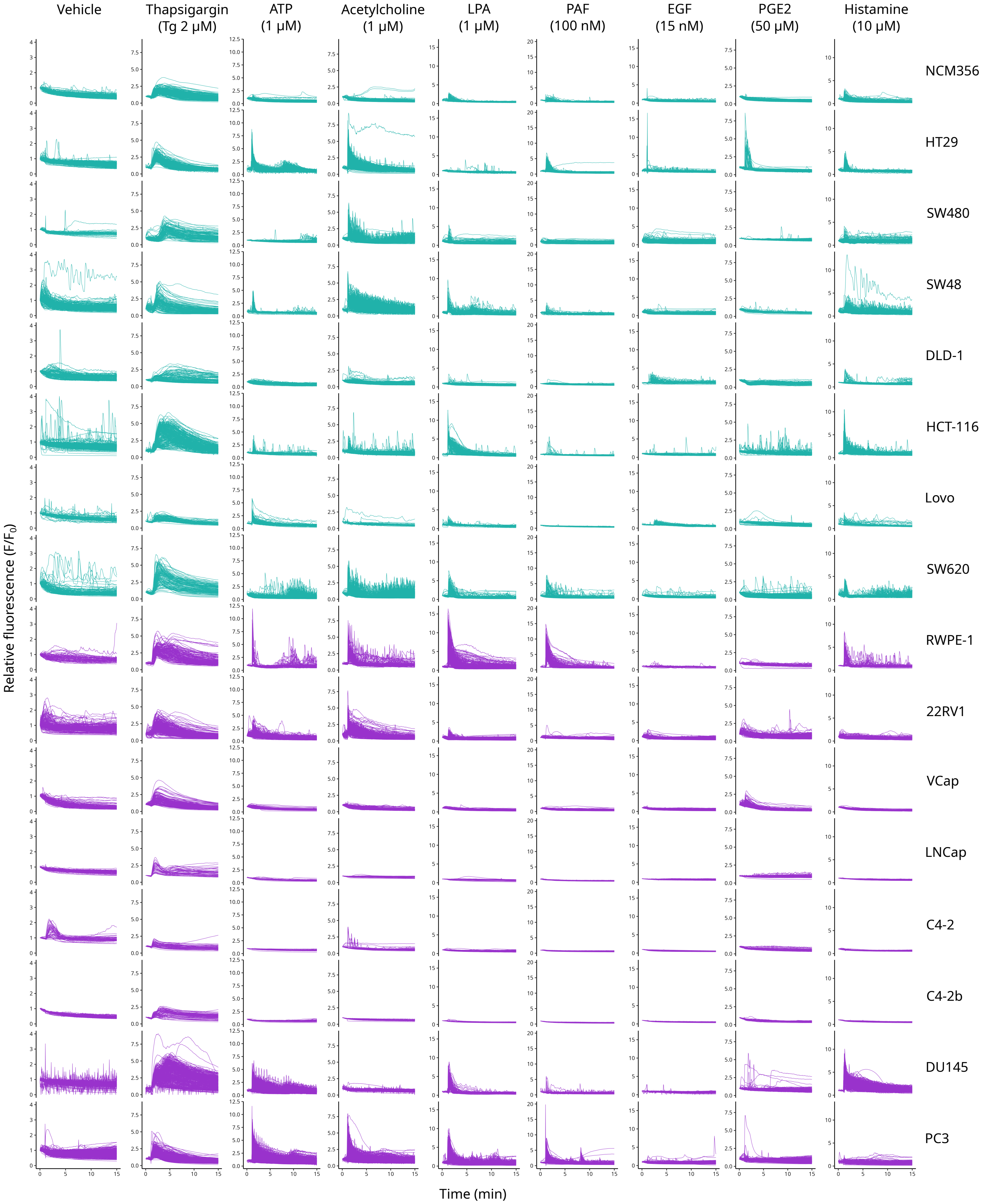

### SUPPLEMENTARY FIGURE 3.

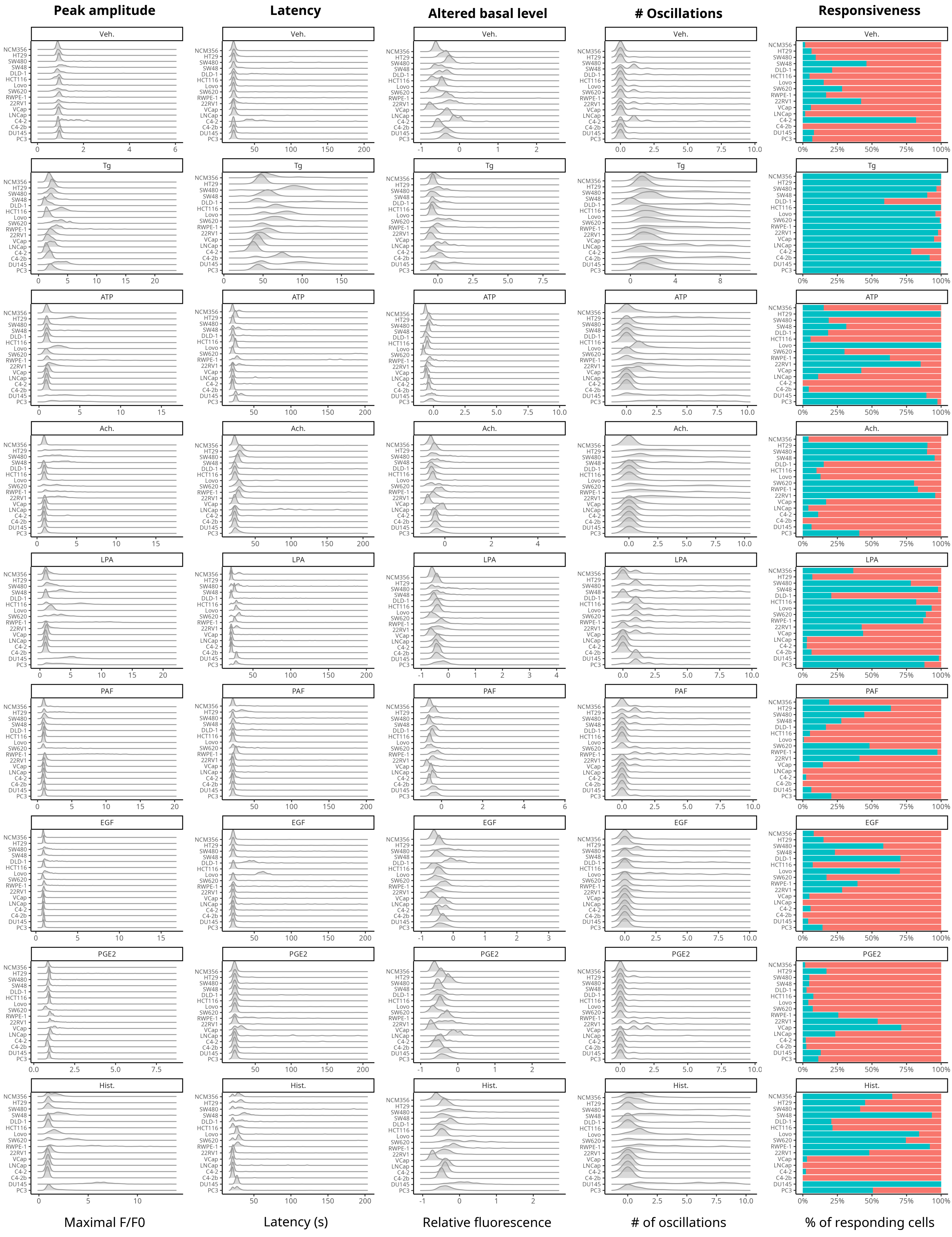

### SUPPLEMENTARY FIGURE 4.

**a.**

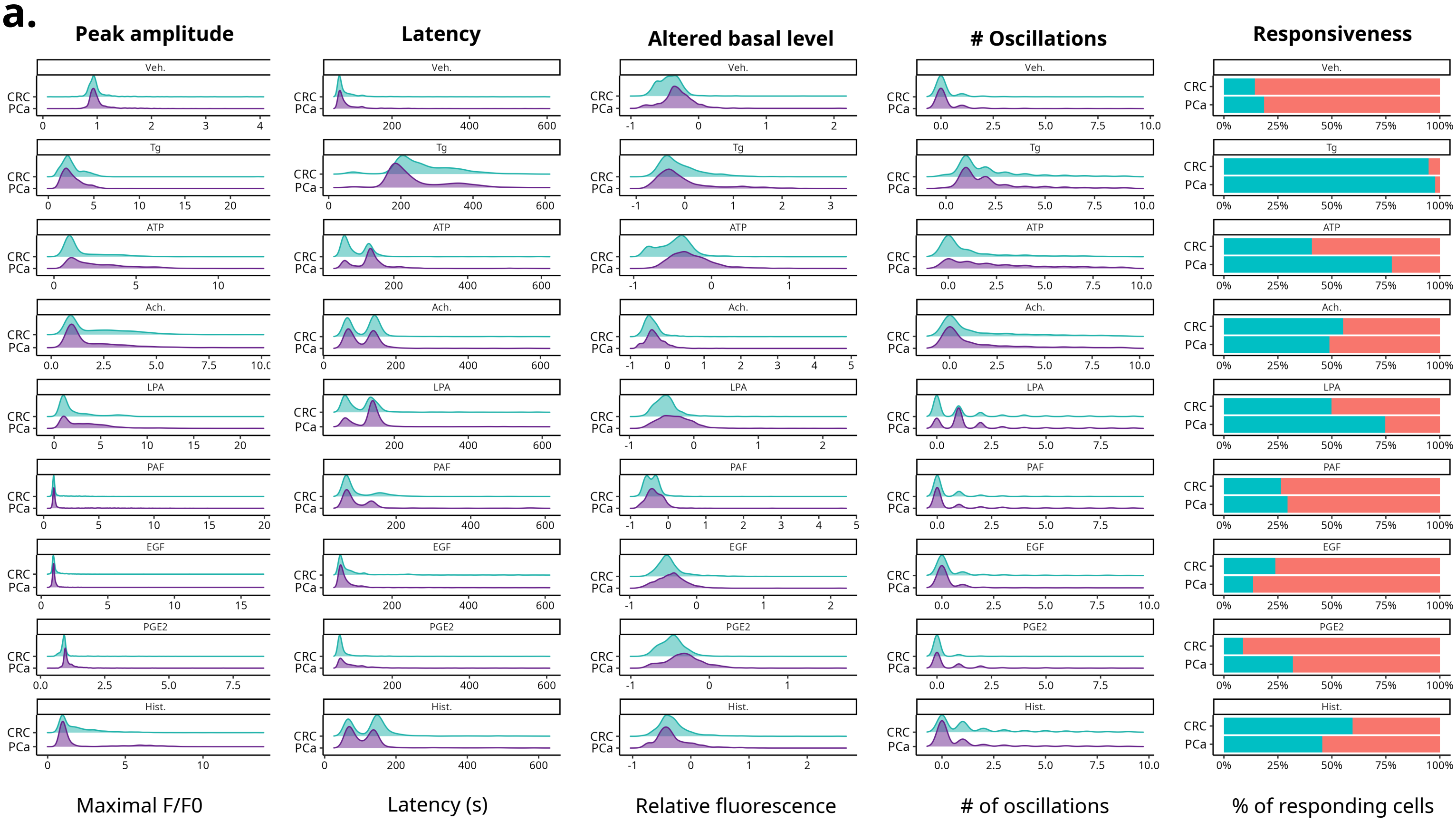


**b.**

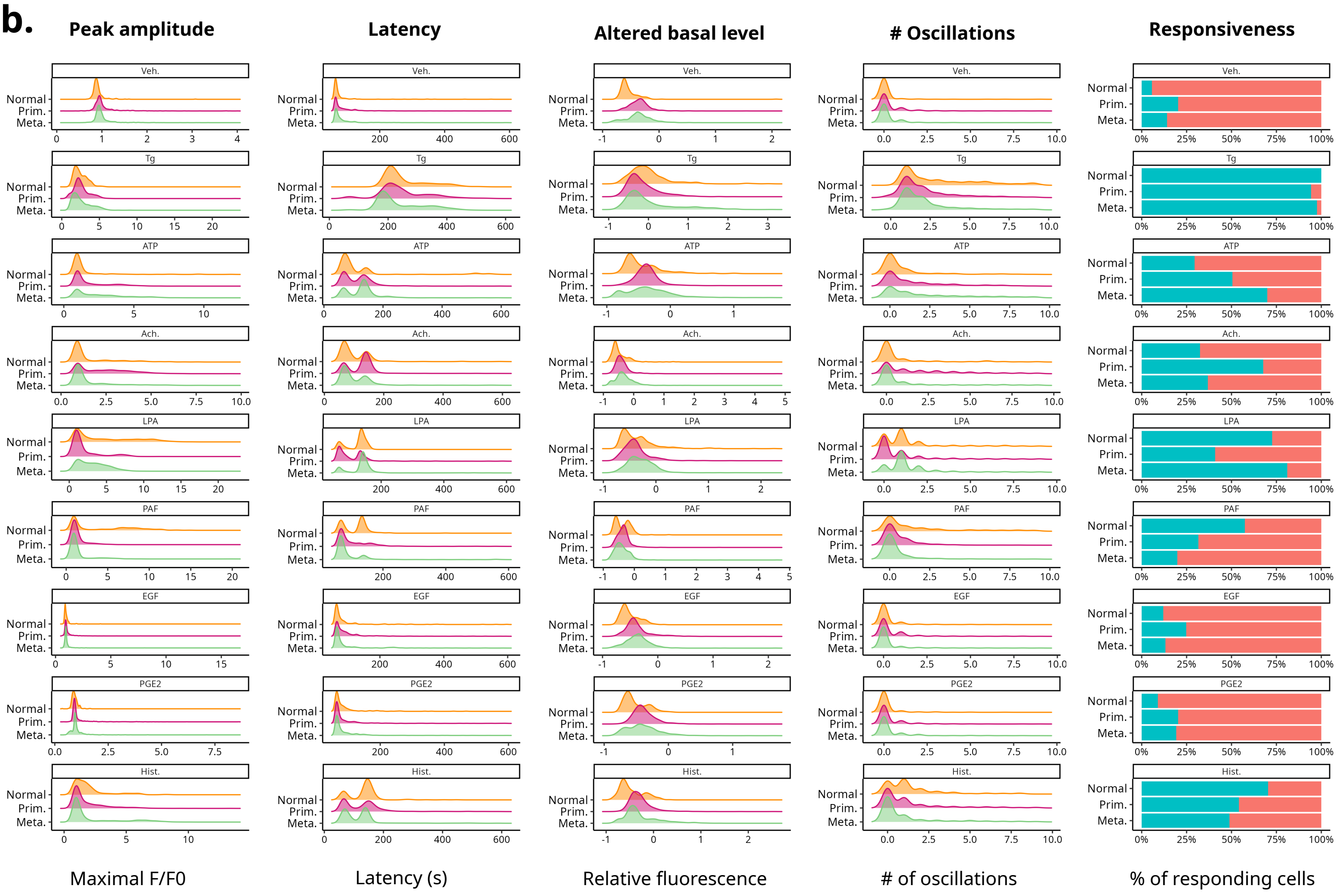

### SUPPLEMENTARY FIGURE 5.

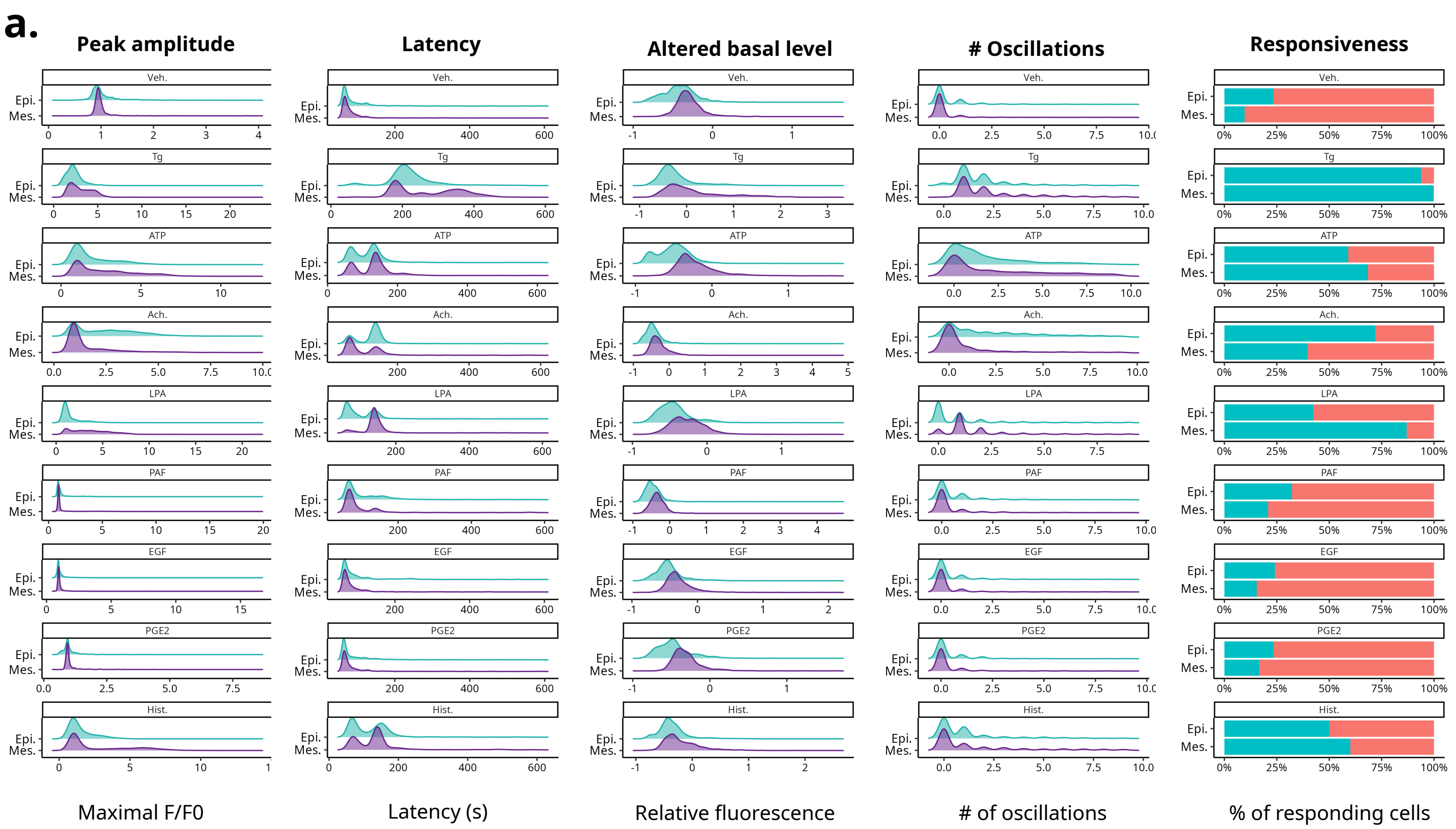

#### SUPPLEMENTARY FIGURE 6.

**a**

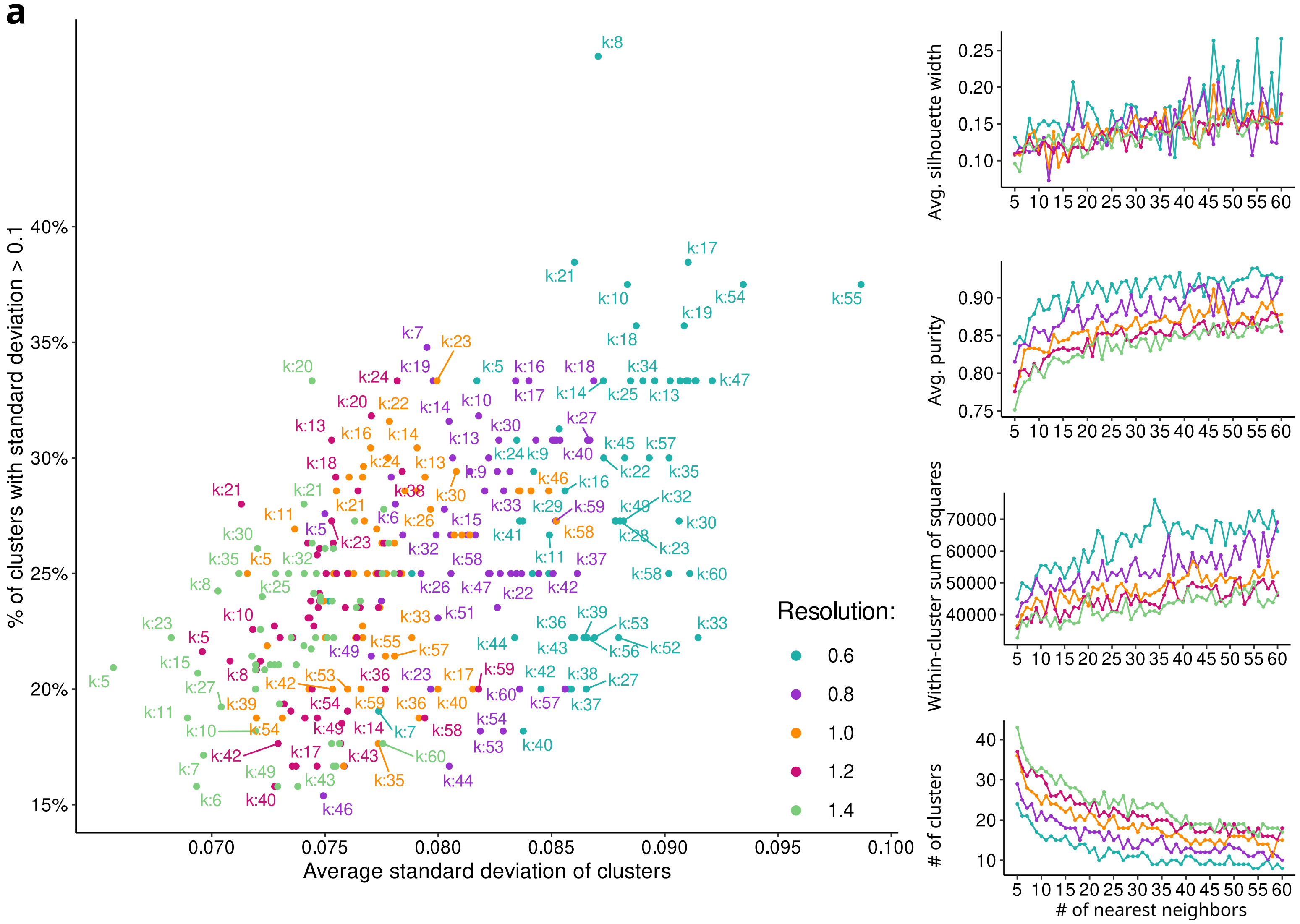

### SUPPLEMENTARY FIGURE 7.

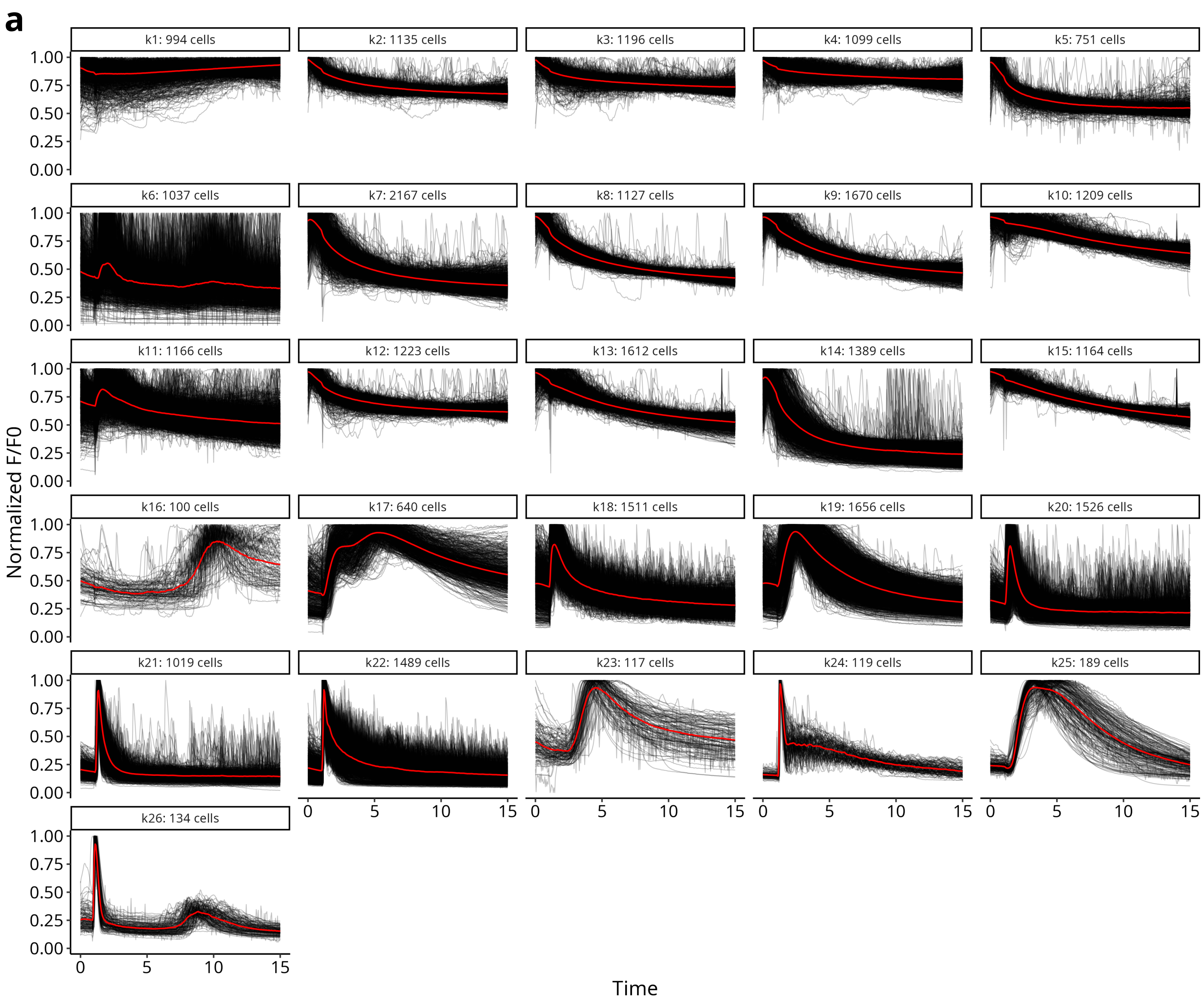

### SUPPLEMENTARY FIGURE 8.

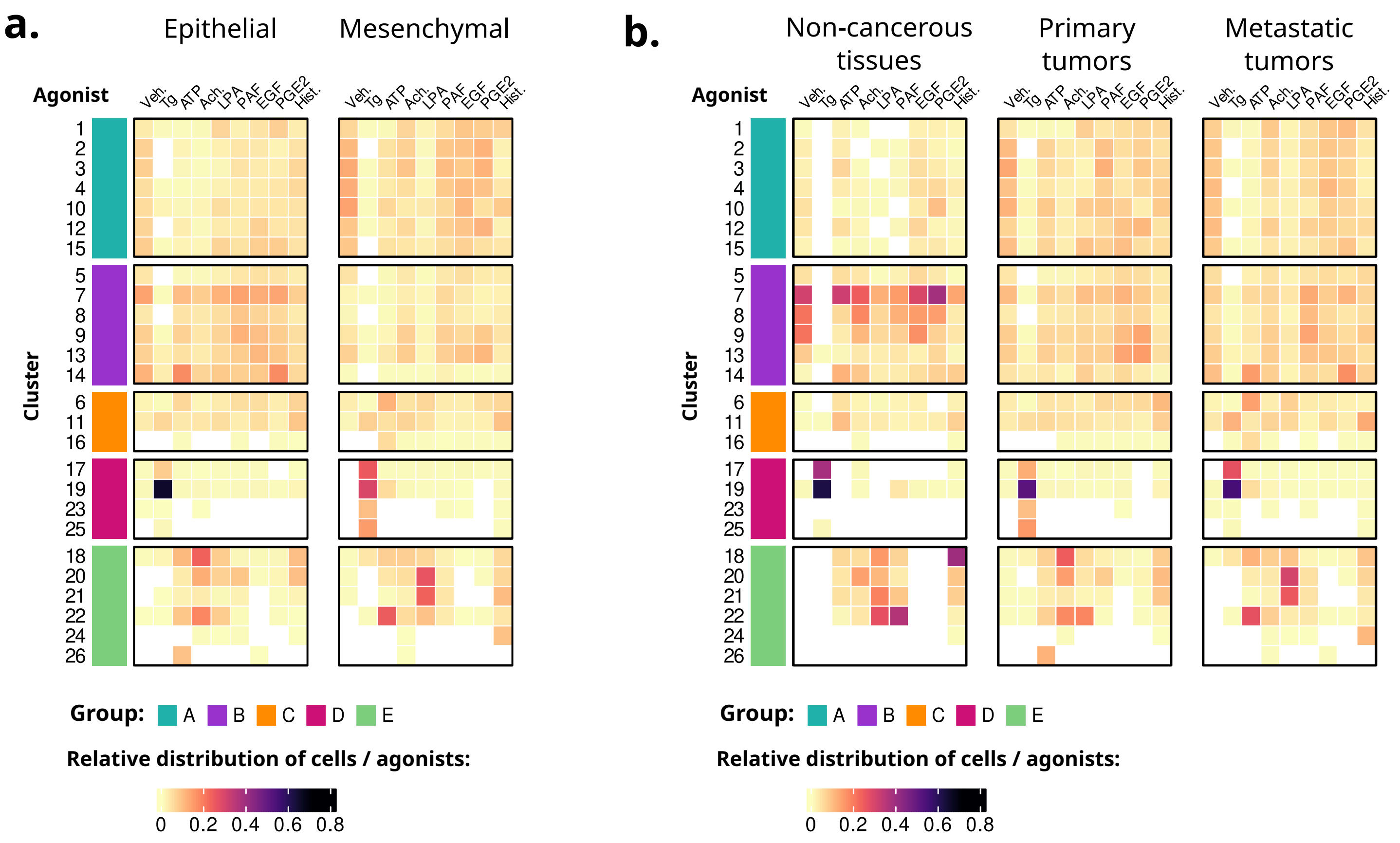

### SUPPLEMENTARY FIGURE 9.

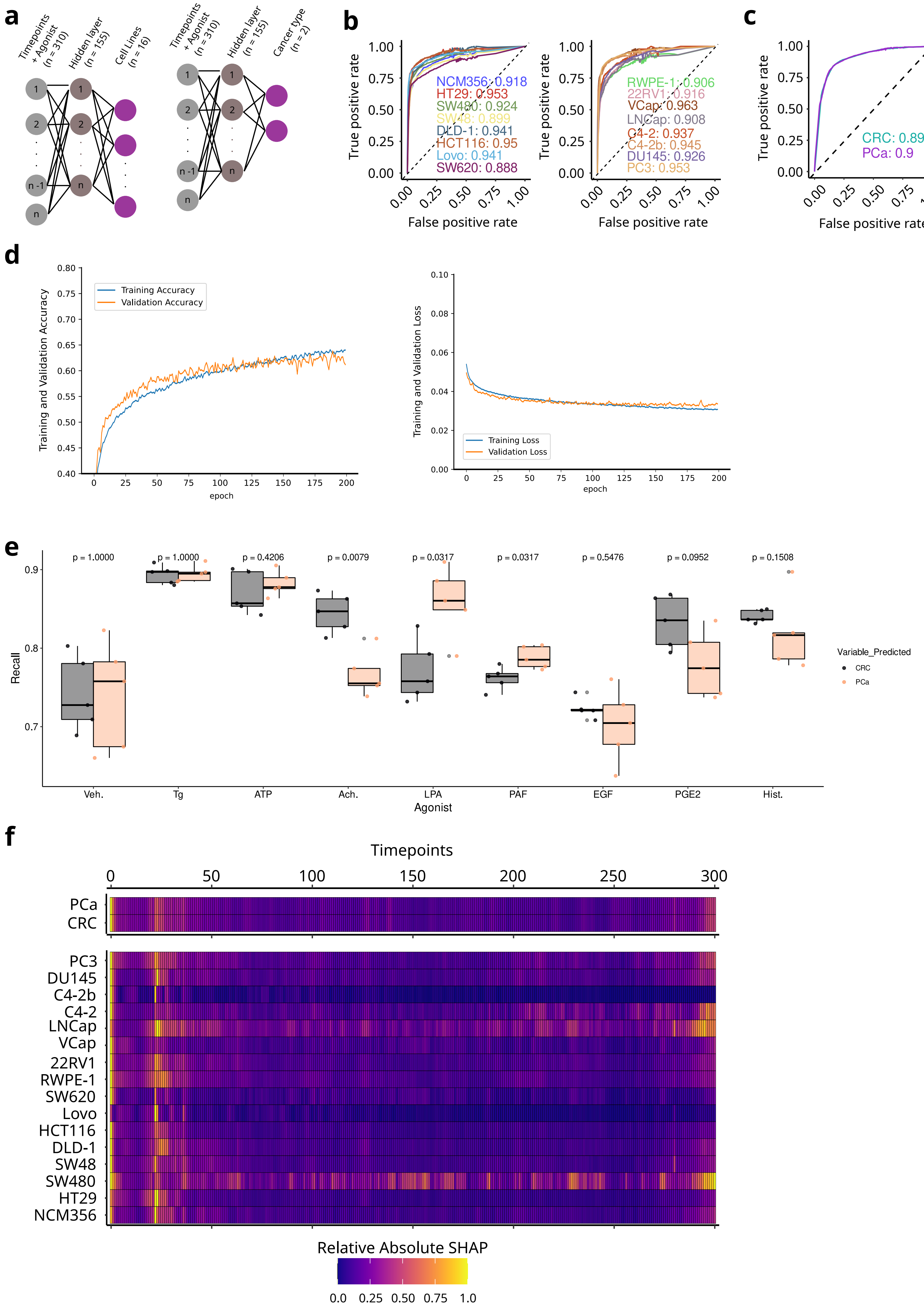

### SUPPLEMENTARY FIGURE 10.

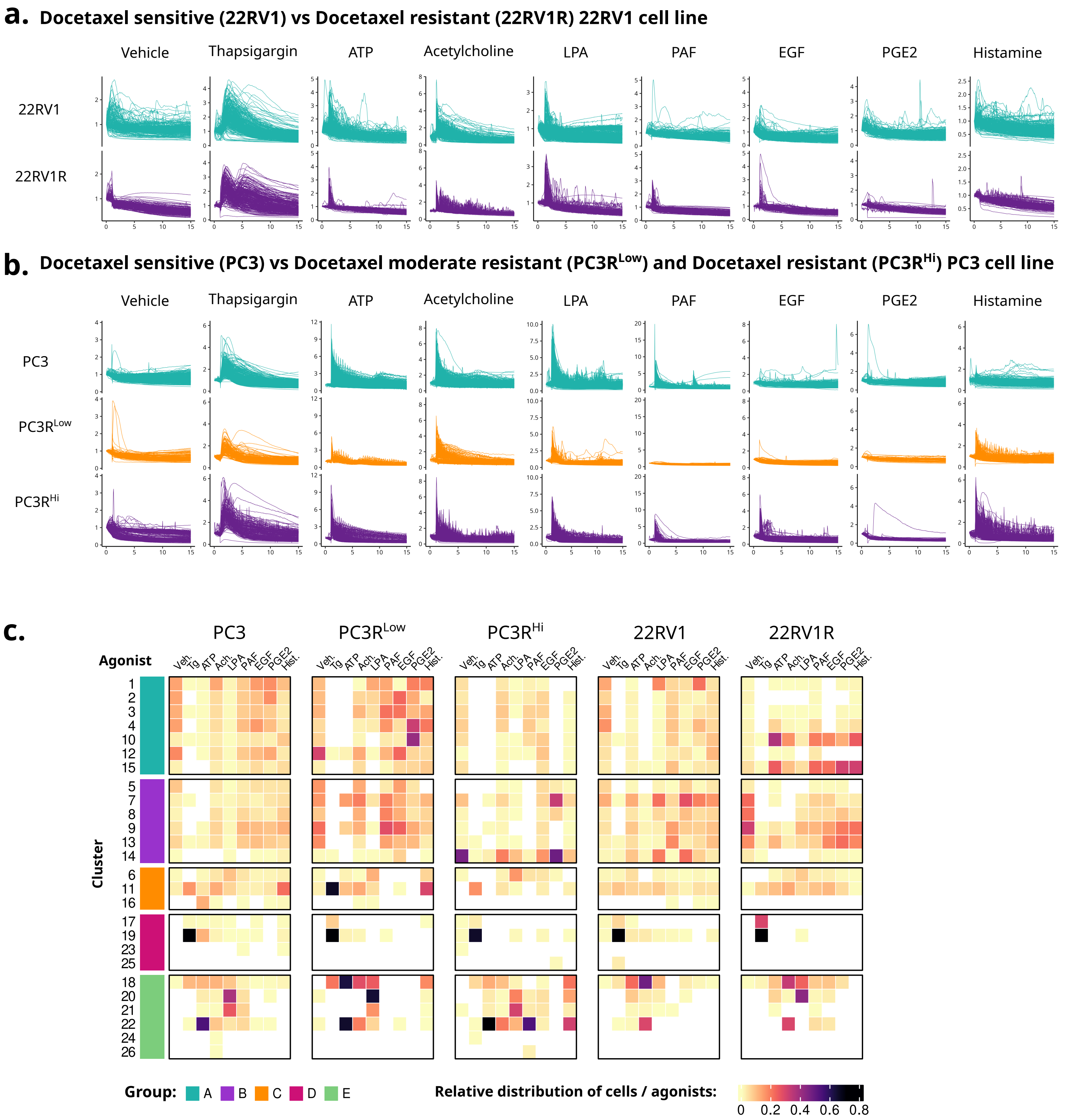

### SUPPLEMENTARY FIGURE 11.

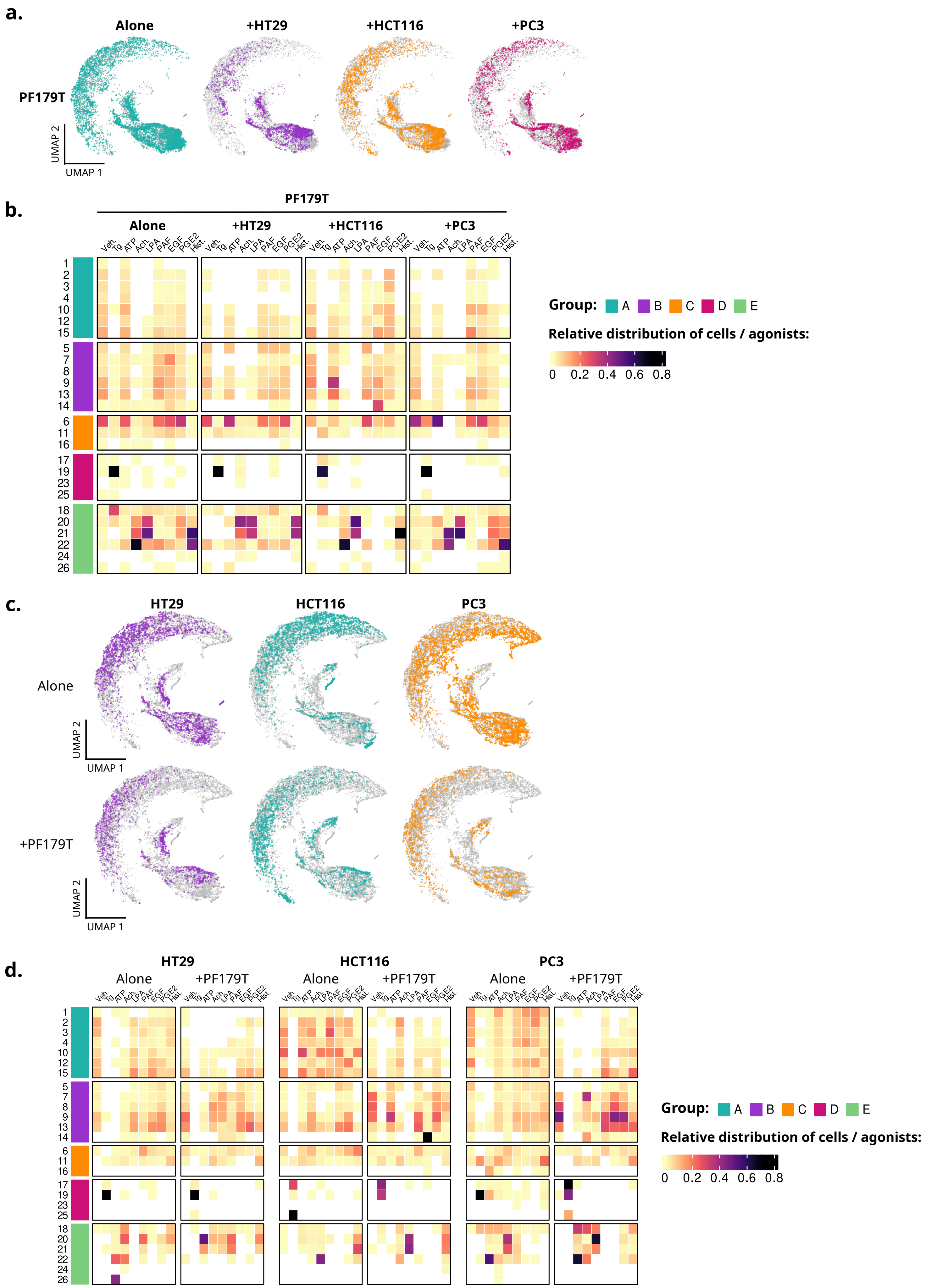
